# An unbiased map of FcγR signaling in neutrophils reveals druggable nodes in immune complex-mediated autoimmune disease

**DOI:** 10.64898/2026.07.21.738176

**Authors:** Mareile Schlotfeldt, Colin Osterloh, Anika Kasprick, Carla Zünkeler, Mikko Armbrust, Georg Kaiser, Katharina Schulze Dieckhoff, Thorge Mester, Pia Stüssel, Nancy Ernst, Ann-Kathrin Schneider, Leonie Voss, Adrian P. Mansini, Seyed Mohammad Vahabi, Yulu Wang, Haley Gainer, Jing Li, Gestur Vidarsson, Remco Visser, Frank Petersen, Xinhua Yu, Kyle T. Amber, Anja Lux, Ralf J. Ludwig, Katja Bieber

## Abstract

Immune complexes drive tissue damage in autoimmune diseases through Fragment crystallizable γ receptor (FcγR) signaling, yet the underlying kinase networks are largely unexplored and treatment remains broadly immunosuppressive. We mapped signaling in immune complex-stimulated human neutrophils by combining an unbiased screen of 155 signal transduction inhibitors with multiplex kinase activity profiling, identifying 51 modulated kinases, 44 of them not previously linked to pemphigoid disease. Nine inhibitors, targeting PLC, PDK-1, PKC, DNA-PK, KSP, BTK, p38, c-Met and TBK-1, reduced neutrophil activity *in vitro* and disease severity in experimental epidermolysis bullosa acquisita, six of them after topical application. BTK emerged as a dominant node as inhibition with ibrutinib was effective therapeutically in immunization-induced disease, in mice reconstituted with a human immune system, and in models of arthritis and immune thrombocytopenia, the latter largely independent of neutrophils. These data provide a resource for FcγR signaling in neutrophils and identify BTK as a cross-disease therapeutic target in antibody-mediated autoimmune disease.

## Introduction

Protein kinases are key players in many essential cellular processes. For example, cell-cycle, proliferation, differentiation, and apoptosis would not be possible without kinases ^1–3^. Protein kinases receive signals from the extracellular environment via different receptors like cytokine receptors, B cell receptors, or fragment crystallizable receptors (FcRs), and transmit the signal to the intracellular space ^1,4^. Here, the signal is amplified triggering downstream events, such as modification of enzymes, gene transcription, or protein degradation ^1,4^. These signaling events occur within interconnected pathways that form complex kinase networks ^5^.

Protein kinases catalyze the transfer of the γ-phosphate from a purine nucleotide (adenosine or guanosine) triphosphate to the hydroxyl group of a protein substrate ^4^. This phosphorylation is a post-translational modification which can be reversed by protein phosphatases. Thereby, the functional state of proteins can be regulated ^4,5^. Protein kinase activity itself is also controlled by phosphorylation which induces conformational changes, thereby activating or inactivating the kinase ^6^. Dysregulation of kinase networks can have far-reaching consequences that can be detrimental for the health of an individual ^7,8^. Consequently, the inhibition of specific kinases has been a rational basis for drug development.

As of July 2026, 100 kinase inhibitors have been approved worldwide and numerous clinical trials are being carried out according to the Protein Kinase Inhibitor Database (PKIDB, http://www.icoa.fr/pkidb)^9^. The success rate of small molecule kinase inhibitors in clinical trials is higher than in general.

Approximately 41 % of small molecule kinase inhibitors are transferred from phase I to approval compared to 18 % of all drugs ^6^. However, only about 30 % of the kinome has been therapeutically targeted so far ^6^, opening the opportunity for further exploration. The screening of small-molecule compound libraries represents a valuable approach for overcoming this treatment gap ^10^.

Most of the inhibitors that were developed up until now have been used in cancer therapy. In contrast, immunological diseases are underrepresented with only 11 small-molecule kinase inhibitors being approved ^6,9^. Importantly, it has been shown that treatments cannot always be transferred from cancer to immunological diseases ^4^. Therefore, more research on targeting signal transduction in immunological diseases is required.

Special examples of immunological disorders are autoantibody-mediated diseases. These autoimmune diseases are characterized by the activation of FcγRs by immune complexes (ICs), i.e. complexes containing immunoglobulin G (IgG) autoantibodies and their antigen, triggering unwanted immune reactions leading to tissue damage ^11^. Prototypical diseases include epidermolysis bullosa acquisita (EBA), rheumatoid arthritis (RA), and immune thrombocytopenia (ITP). Well-established animal models exist for each ^12–14^, and at least RA and pemphigoid diseases (PDs) such as EBA ^15^ require more targeted treatment options.

Importantly, this shared pathogenic mechanism is executed by distinct FcγR-expressing effector cell populations, which may differentially engage intracellular signaling pathways. Although neutrophils are among the most prominent effector cells in experimental EBA ^16,17^, the infiltrates in human patients may also contain eosinophils and lymphocytes in the papillary/superficial dermis ^18^. Important characteristics of neutrophil activation are chemotaxis and the release of reactive oxygen species (ROS) release, and cytokines ^19^, which are responsible for tissue damage ^17,20^. In RA, besides neutrophils, macrophages and mast cells, among others, also play an important role ^21–23^. In contrast, neutrophils are only minorly involved in ITP ^24,25^. Moreover, even though kinases are ubiquitously expressed among cell types, their activation and downstream signaling can differ substantially. These differences may be explained by cell type-specific receptor expression and kinase phosphorylation, as shown for Bruton’s tyrosine kinase (BTK) ^10^.

A small number of kinase inhibitors have been successfully tested in pre-clinical models of these three diseases. Several JAK inhibitors are approved for RA and the SYK inhibitor fostamatinib is in clinical use for ITP ^9^. Yet, no larger screens have been conducted, and the signaling networks remain largely unknown. Therefore, this study set out to analyze how signal transduction can be targeted in these disease models, first using EBA as a proof-of-concept model and then extending the analysis to models of RA and ITP. By screening of a target-selective library containing small-molecule signal transduction inhibitors (STIs) and multiplex kinase activity profiling of human IC-stimulated neutrophils, we identified potential drug targets for IC-mediated autoimmune diseases. We validated these candidates *in vitro* and *in vivo* using different models for EBA, RA, and ITP. Regarding EBA, both prophylactic and therapeutic interventions as well as a recently developed humanized EBA model were performed. Thus, the present study provides a major addition to the elucidation of the signaling pathways in immune complex-activated neutrophils. Furthermore, we contribute to potentially enhanced treatment options for patients with neutrophil-driven FcγR-dependent autoimmune diseases in the future.

## Results

### Identification of 59 signaling molecules as potential new drug targets in IC-mediated autoimmune diseases

To identify druggable signaling molecules in IC-activated neutrophils, we combined two complementary approaches. To analyze which kinases are modulated in human IC-stimulated neutrophils, a multiplex kinase activity profiling was performed. Additionally, we screened a target-selective compound library comprising 155 STIs targeting 151 different signaling molecules in a ROS release assay using human IC-stimulated neutrophils. Based on this dual screening approach, we selected several inhibitors for both *in vitro* and *in vivo* validation (Fig. 1) as described in the following paragraphs.

**Fig. 1:**
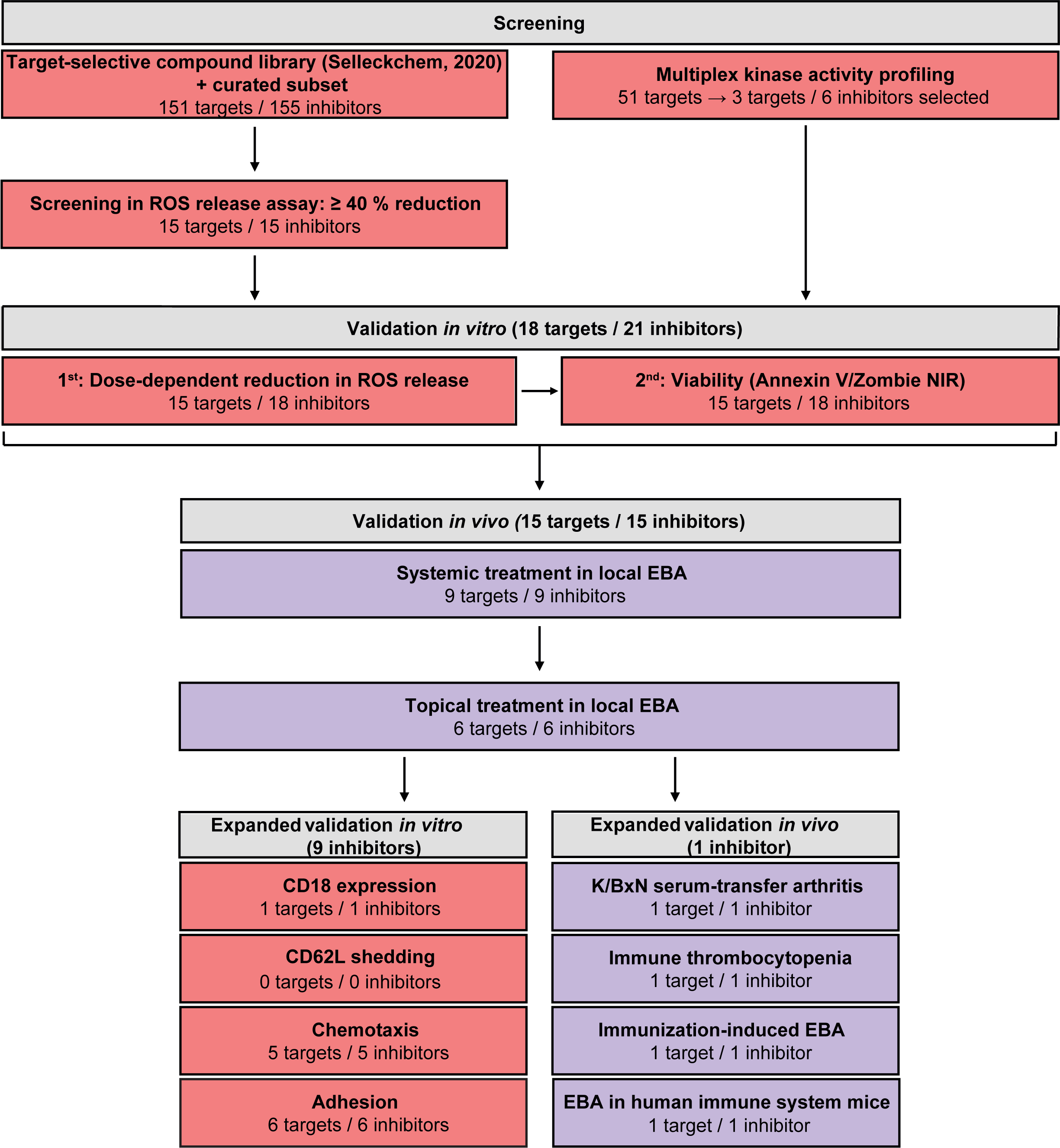
Screening and validation of signal transduction inhibitors as targets in immune complex-induced autoimmunity. Following an initial screen of a target-selective inhibitor library in a ROS release assay and a multiplex kinase activity profiling, both conducted with IC-stimulated human neutrophils, 23 inhibitors were validated *in vitro.* If they dose-dependently reduced ROS release and showed no toxic effects, they were applied in experimental EBA *in vivo.* The nine inhibitors that were effective *in vivo* were selected for expanded validation *in vitro,* assessing CD18 expression, CD62L shedding, chemotaxis and adhesion. The most effective inhibitor, ibrutinib, was chosen for expanded validation *in vivo* in K/BxN STA, ITP, immunization-induced EBA, and EBA in HIS mice. *In vitro* experiments are represented in red, *in vivo* experiments in blue.

The multiplex kinase activity profiling confirmed the modulated activity of 51 kinases in IC-stimulated neutrophils compared to unstimulated neutrophils (Fig. 2a), out of which seven kinases were already suggested to be implicated in EBA, including ERK1/2 ^26^, p38 ^26^, BTK ^27^, CDK7, CDK9 ^28^ and ERK5 ^29^. Moreover, we detected the modulated activity of multiple kinases that had not been previously described in EBA, such as ROCK1/2, NUAK1, PAK1 and many more (Supplementary table 3). However, selective STIs were not available for all targets. Consequently, we chose three targets with sufficiently selective inhibitors available for further investigation: TBK1, DDR1 and BTK. These three kinases were further validated as potential therapeutic targets. Given that BTK inhibition by ibrutinib has already been described as an effective treatment option for antibody transfer-induced EBA ^27^, we selected three additional STIs for optimization and validation of previous results.

**Fig. 2:**
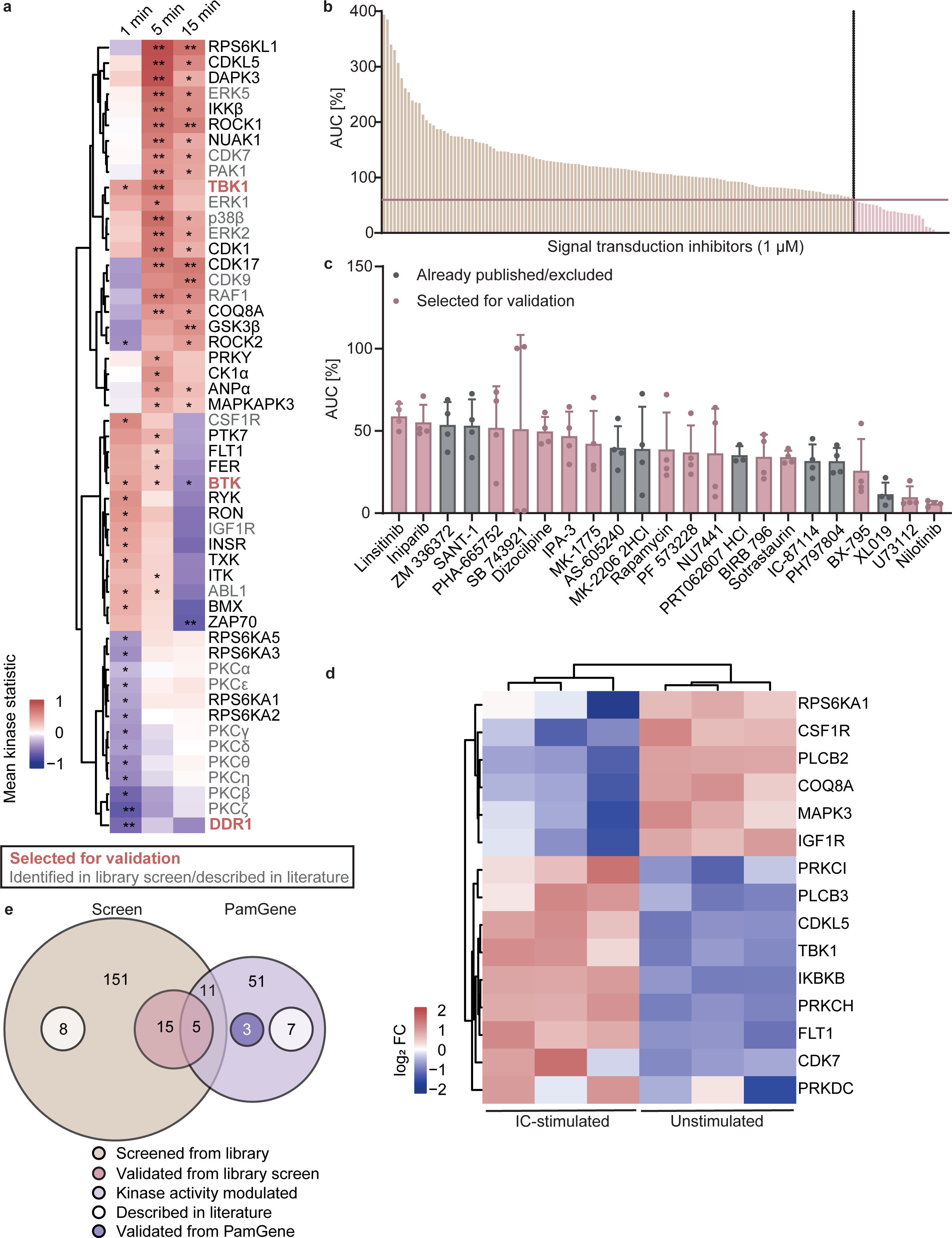
Multiplex kinase activity profiling and compound library screening reveal potential novel therapeutic targets for IC-mediated autoimmune diseases. Human neutrophils were isolated and stimulated with immobilized IC consisting of human COL7 E-F and anti-human COL7 IgG1 (COL7-IC). (a) Lysates were prepared after 1 min, 5 min, and 15 min of stimulation, and analyzed by multiplex kinase activity profiling using PamGene™. * mean specificity score ≥ 1, ** mean specificity score ≥ 1 and mean significance score ≥ 0.5. (b) A curated target-selective library of signal transduction inhibitors (STIs) was screened at 1 µM for reduction of ROS release by at least 40 % after IC stimulation for 2 h. The mean for each inhibitor is shown, individual values can be found in Supplementary tables 2 and 3. **(c)** STIs below this threshold are shown in detail. Data are represented as scatter dot plots (mean + SD) **(d)** Human neutrophils were isolated by MACS, stimulated with immobilized COL7-IC for 6 h and prepared for bulk RNAseq together with unstimulated neutrophils from the same donor. Significantly differentially expressed genes (p ≤ 0.1) were identified and log2 fold changes (log2 FC) of kinases identified in multiplex kinases activity profiling are shown. **(e)** A Euler diagram shows the overlap between the PamGene experiment and the library screen and indicates whether the STIs were chosen for validation in the local antibody transfer-induced EBA model. **(a, d)** n = 3. **(b, c)** n = 4.

In addition, a library of 155 STIs was screened using a ROS release assay of IC-activated human neutrophils (Fig. 2b, Supplementary table 1). This resulted in eleven overlapping targets with the kinase activity measurement, six of which passed the validation threshold of at least 40 % reduction in ROS release. Considering the largely unbiased screening approach, we deem this a substantial number of overlaps. A total of 15 inhibitors reducing ROS release by at least 40 % were selected for further validation, excluding compounds targeting molecules already implicated in EBA, namely PI3K ^30–32^, AKT1 ^26^, SYK ^33,34^, CDK7 ^28^, ERK2 ^26^, and SRC ^35^ (Fig. 2c). BIRB 796 targeting p38, which has previously been identified as involved in EBA ^26^, was chosen as positive control to strengthen our results. Two inhibitors (ZM 336372 and SANT-1) were excluded since to our knowledge, no data on tolerability *in vivo* have been published. This results in 59 novel targets in total have been identified by at least one of the two approaches. Out of these targets, 18 targets and 21 inhibitors were selected for validation (Fig. 2e, Supplementary table 29).

Since kinase activity can also be regulated by alteration of gene expression, a bulk RNAseq experiment was performed using human neutrophils, either unstimulated or stimulated with IC for 6 h (Fig. 2d, Fig. S1). Out of the targets selected for validation, the analysis revealed the significant upregulation of several *PRKC* isoforms, *PRKDC*, *PLCB2*, and *IGFR1*. Together, these findings highlight the probable importance of 18 novel signaling molecules identified by both library screening and multiplex kinase activity profiling in neutrophil signaling in IC-mediated diseases, such as EBA (Supplementary table 29).

### ROS release is reduced in a dose-dependent manner by 18 signal transduction inhibitors

Since ROS release is a key effector function of neutrophils and responsible for tissue damage in neutrophil-mediated diseases including EBA ^17^, a dose-dependent ROS release assay with four inhibitor concentrations ranging from 0.01 µM to 10 µM was performed for further analysis of the inhibitors selected from Fig. 2a and for validation of those identified in Fig. 2c. Lower concentrations were chosen if a reduction, but no dose dependency was observed at these concentrations. We found 18 out of 23 tested substances to dose-dependently reduce ROS release from human IC-stimulated neutrophils: U73122, BX-795, sotrastaurin, BIRB 796, NU7441, SB 743921 HCl, PHA-665752, GSK8612, ibrutinib, PF 573228, rapamycin, MK-1775, linsitinib, DDR1-IN-1, nilotinib, CC-292, orelabrutinib, and BMS-986142 (Fig. 3a, Fig. S2a). To exclude that these effects were simply due to toxicity of the compounds, a viability test was conducted for those STIs that reduced ROS release (Fig. 3b, Fig. S2b). Significant toxicity was not observed for any of the STIs except for U73122 and SB743921 HCl at 10 µM. However, for PHA-665752 and sotrastaurin compounds the evaluation at high concentrations was impossible due their intrinsic color, which interfered with the flow cytometric assay. Therefore, a trypan blue staining was conducted showing that viability of the cells was not affected (Fig. S2c). Hereby, we confirmed the potential of 18 inhibitors targeting 15 signaling molecules to decrease ROS release from human neutrophils.

**Fig. 3:**
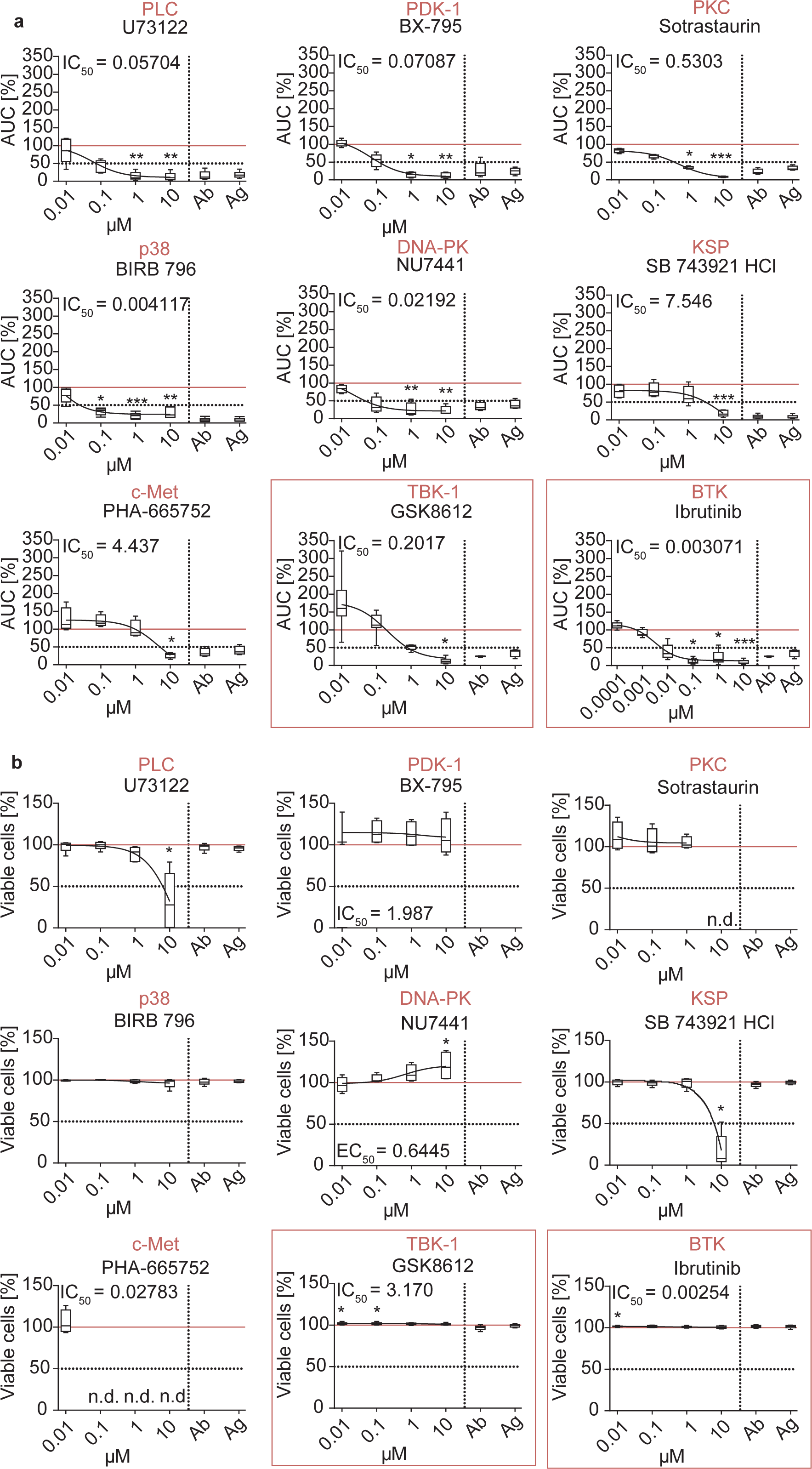
Validation of nine STIs in a dose-dependent ROS release assay and viability measurement *in vitro*. Human neutrophils were purified and treated with immobilized human COL7-IC and STIs for 2 h. Values are normalized to the positive control (IC-stimulated neutrophils treated with DMSO/DMF only). Cells treated with antibody (ab)-or antigen (ag) only served as negative controls. **(a)** ROS release was continuously measured in a luminol-based assay. **(b)** The cells were stained with Annexin V and Zombie NIR to assess viability. The data are represented as Tukey box-and-whisker plots. The red horizontal line indicates the positive control, which was used for normalization. IC50 curves are depicted and IC50 values are stated if within the tested concentration range. n.d., not determined. Kruskal-Wallis test with Dunn’s post-test was performed. * p ≤ 0.05. ** p ≤ 0.01. *** p ≤ 0.001. n = 5. n.d., not determined.

### Prophylactic treatment with nine systemically applied STIs diminishes local EBA

To analyze the effect of pharmacological blockade of these 15 signaling molecules on local antibody transfer-induced EBA in mice, STIs were systemically administered starting before EBA induction with anti-murine collagen VII (mCOL7)^c^ IgG (Fig. 4a). The mice were observed for a total of five days. The results are exemplified by clinical images, hematoxylin and eosin (H&E) stainings and IgG and C3 deposition from BX-795-treated mice and controls (Fig. 4b). Nine inhibitors (U73122, BX-795, sotrastaurin, BIRB 796, NU7441, SB 743921 HCl, PHA-665752, GSK8612, and ibrutinib) reduced clinical disease manifestation, i.e. the affected ear surface area (AESA) and/or the difference in ear thickness (Fig. 4c). This was frequently accompanied by histological changes (Fig. S4). Reduced inflammation was observed in H&E stainings of the mice treated with BX-795, BIRB 796, SB 743921 HCl, PHA-665752, and ibrutinib. Notably, the latter mitigated all three tested parameters: epidermal thickness, cell infiltration, and split formation. In contrast, six inhibitors (DDR1-IN-1, nilotinib, PF 57228, rapamycin, MK-1775, and linsitinib) had no effect on either difference in ear thickness nor AESA (Fig. S3).

**Fig. 4:**
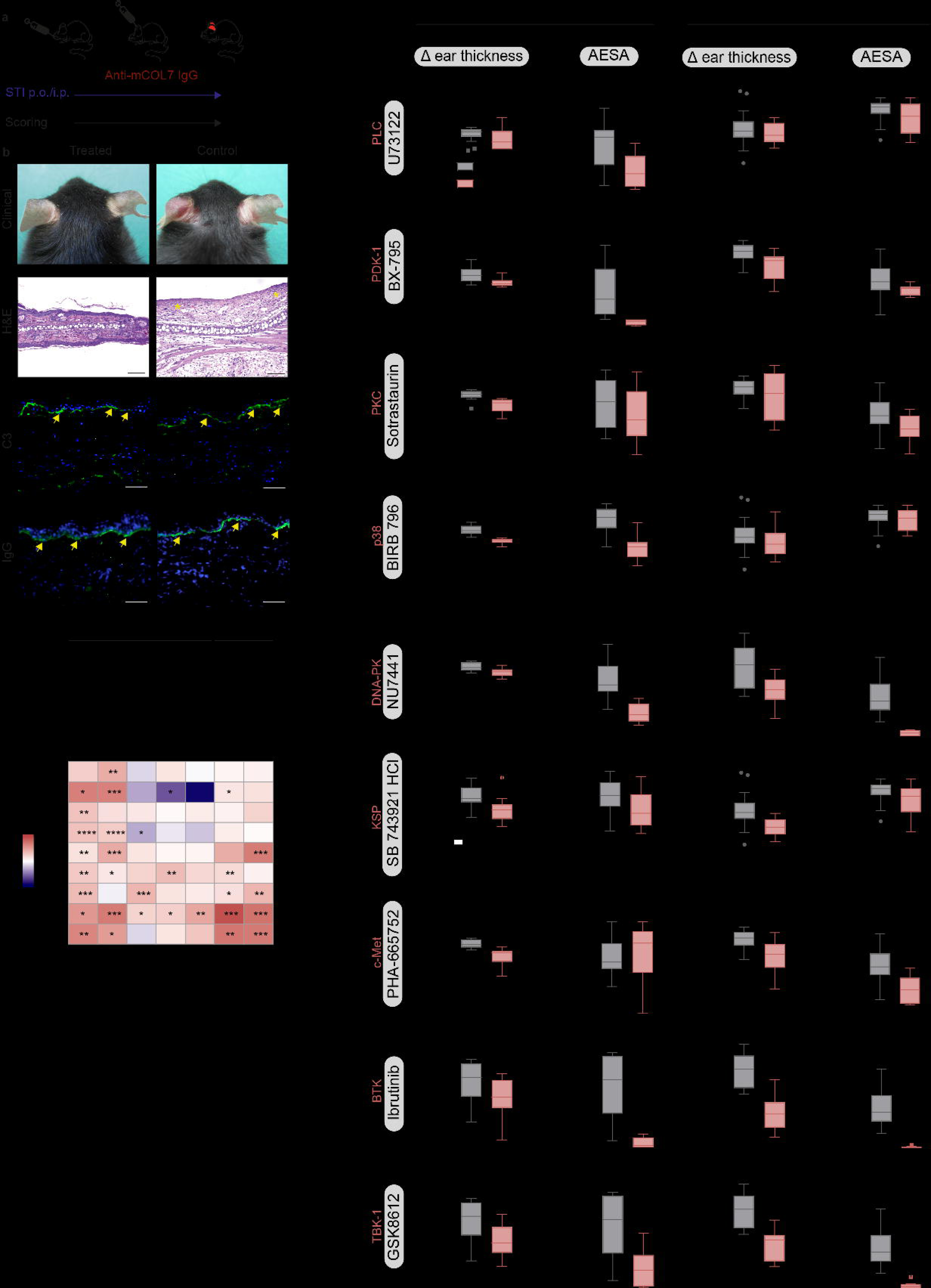
Nine STIs are confirmed as potential therapeutic options in the local antibody transfer-induced EBA model. **(a)** Mice were treated with STIs before the induction of local antibody transfer-induced EBA on day 0 and clinically scored every day for five days. **(b)** Representative clinical images, H&E, C3, and IgG stainings from BX-795-treated and control mice are shown. Asterisks indicate cell infiltrates and arrows indicate binding of C3 or IgG to the dermal-epidermal junction. **(c)** Upon systemic STI treatment, the affected ear surface area (AESA) and the difference in ear thickness were determined on the final day of the experiment. **(d)** Analogous representation for topical STI treatment. **(e)** Overview of the reduction in the respective parameter comparing the means of each treatment and control group including histological data for the systemic treatment. Mann-Whitney test was performed. * p ≤ 0.05. ** p ≤ 0.01. *** p ≤ 0.001. n = 4 (ibrutinib, GSK8612), n = 5 (BX-795, sotrastaurin, NU7441, PHA-665752), n = 8 (U73122, BIRB 796, SB 743921 HCl).

Topical treatment could limit systemic exposure and thereby reduce the risk of adverse effects, a potential advantage for skin disease. We therefore tested the nine *in vivo*-validated STIs topically in the same model (Fig. 4d). Here, six out of nine STIs (BX-795, NU7441, SB 743921 HCl, PHA-665752, ibrutinib, and GSK8612) reduced the AESA and/or the difference in ear thickness. Thus, prophylactic application of selected STIs, especially ibrutinib, can improve local EBA (Fig. 4e).

### Further *in vitro* functional validation unveils effects on surface activation marker expression, chemotaxis, and adhesion

For a more detailed mechanistic insight into the neutrophil functions that are impaired by the STIs that were successfully used in local EBA, additional *in vitro* functional assays were performed. Shedding of CD62L remained unchanged for all STIs, except for GSK8612, which led to an increase of CD62L^-^cells at 0.1 µM. (Fig. S5a). CD18 expression was significantly reduced only by ibrutinib (Fig. S5b). Adhesion was dampened by U73122, BX-795, sotrastaurin, SB 743921 HCl, ibrutinib, and GSK8612 in a dose-dependent manner (Fig. S6a). While all these functions are based on IC-mediated signaling, we addressed the question whether FcγR-independent pathways are also affected by STI treatment. Therefore, an IL-8-dependent chemotaxis assay was conducted, showing reduced migration for U73122, BX-795, SB 743921 HCl, ibrutinib, and GSK8612 (Fig. S6b). However, it should be noted that U73122 and SB 743921 HCl were toxic at 10 µM, suggesting that the observed effects at this concentration may have been caused by a reduced number of viable cells. Thus, ibrutinib demonstrated the greatest capacity to reduce neutrophil activity among the STIs.

### Ibrutinib mitigates EBA in the immunization-induced model and blocks EBA induction in humanized immune system (HIS) mice

To assess the role of BTK in a therapeutic EBA setting, an immunization-induced EBA experiment was performed. Mice were immunized with mCOL7^vWFA^^2^ and after development of EBA, they were randomized into treatment and control groups (Fig. 5a). Over 85% of mice treated with ibrutinib reached clinical remission, whereas none of the control group did (Fig. 5b). The treatment with ibrutinib diminished the disease starting in week 1 and further improved until week 4 (Fig. 5c). This enhancement was further substantiated by the clinical images over the time course of the experiment (Fig. 5d) and on the histological level by H&E staining (Fig. 5e). Accordingly, the quantification of the histological stainings showed a reduction in epithelial thickness and cell infiltration. C3 deposition was decreased in the ibrutinib-treated mice, while IgG deposition was unchanged. (Fig. 5f). Serologically, no difference in total IgG was observed between the two groups, but a lower anti-mCOL7^c^ titer in the control mice. Importantly, none of the individual anti-mCOL7^c^ IgG subclasses were significantly affected, especially not the most pathogenic subclasses IgG2b and IgG2c (Fig. S7), indicating a clear ibrutinib-mediated impairment of the effector phase of inflammation, but not on the induction phase.

**Fig. 5:**
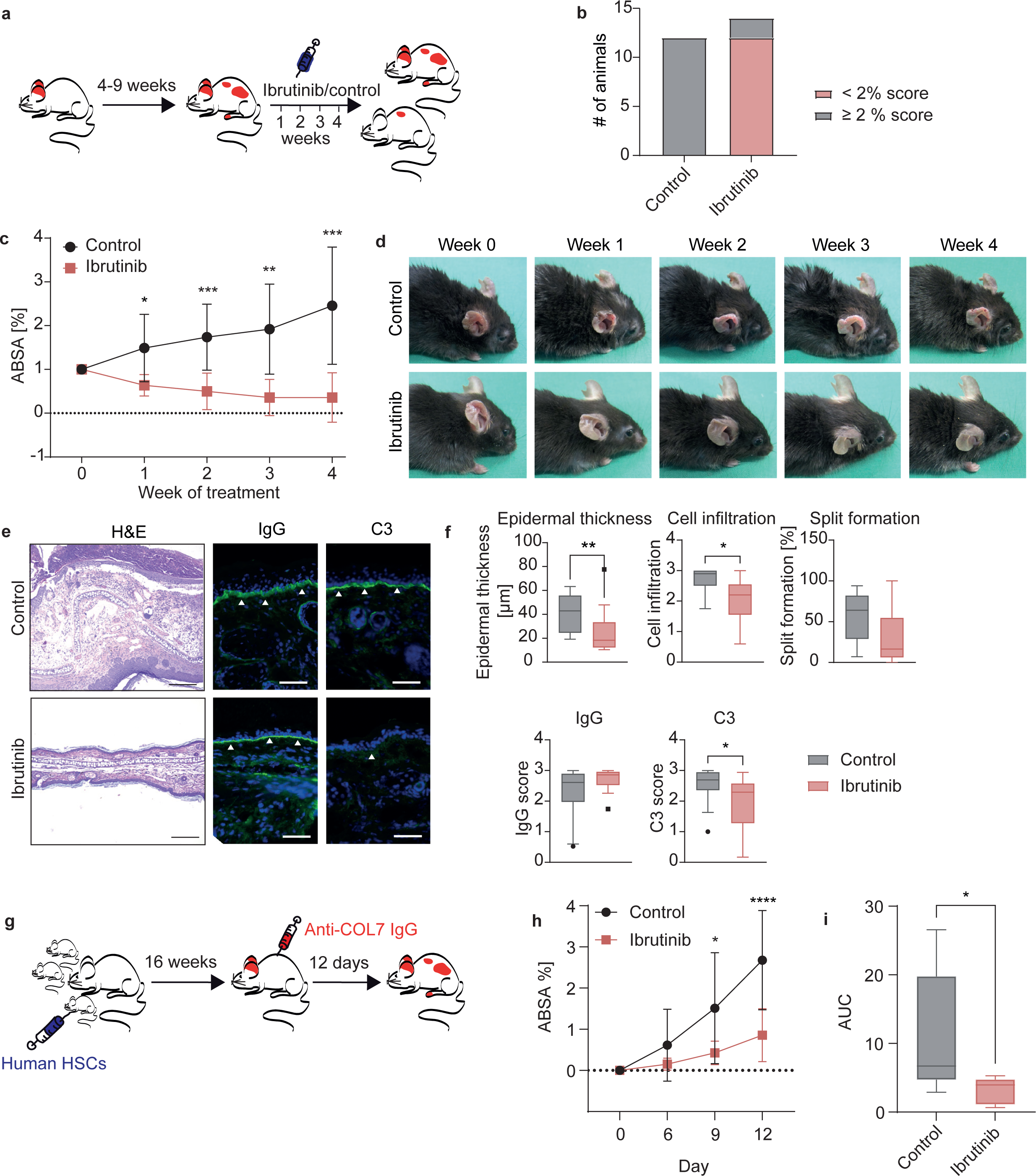
Both active EBA in B6.s mice and passive EBA in humanized immune system mice can be mitigated by ibrutinib. **(a)** Active EBA was induced in B6.s mice and animals that developed an affected body surface area (ABSA) of ≥ 2 % were randomized into the treatment groups. **(b)** Animals with an ABSA of < 2 % were considered in remission and the number of animals below and above this threshold is shown for the two groups. **(c)** Development of the disease score over the four weeks of treatment. n = 12 (control), n = 14 (ibrutinib). Two-way ANOVA with Šidák’s multiple comparisons test was performed. **(d)** Representative clinical images over the time course of an individual mouse. **(e)** Representative images from H&E, IgG, and C3 stainings are shown. Arrows indicate the deposition of IgG or C3. **(f)** Epidermal thickening, cell infiltration, and split formation were quantified. Mann-Whitney test was performed. **(g)** Induction of EBA in the humanized immune system mouse model. Newborn NSG-FcRγ^−/−^ mice were irradiated and injected with human HSCs 16 weeks after transplantation. Then, EBA was induced by anti-COL7 IgG injection and mice were treated for twelve days with ibrutinib or vehicle. **(h)** Development of the affected body area during the treatment. The ABSA was evaluated and two-way ANOVA with Šidák’s multiple comparisons test was performed. **(i)** The area under the curve (AUC) over the 12-day experiment for each group was calculated and Mann-Whitney test was conducted. n = 7 (control), n = 9 (ibrutinib). **(c, f, h, i)** * p ≤ 0.05. ** p ≤ 0.01. *** p ≤ 0.001, **** p ≤ 0.0001.

To improve translational predictability, ibrutinib was also tested in a recently developed ^36^ HIS EBA mouse model. Here, mice were irradiated and then transplanted with HSCs. EBA-like disease was induced by injection of anti-COL7^c^ rabbit IgG and treated with ibrutinib or vehicle for twelve days (Fig. 5g). The disease score was significantly lower in the treated mice on day 9 and the difference became even clearer on day 12 (Fig. 5h). This was also reflected by the area under the curve (AUC, Fig. 5i). At the cellular level, a significant decline in the number of human classical monocytes as well as a slight reduction in human neutrophils was observed in the skin (Fig. S8a), while the number of murine cells remained constant (Fig. S8b). Furthermore, both human (Fig. S9a) and murine cytokine levels (Fig. S9b) were unaffected, except for murine IL-17A which was reduced in the treatment group on day 13. However, the biological relevance of this change is likely negligible due to the minimal concentration. Other cytokines such as IL-6, IL-33, IL-12p70, and IL-18 were expressed at higher concentrations and showed a tendency to decrease in treated mice.

Overall, the efficacy of ibrutinib in these two additional EBA models highlights BTK as promising therapeutic target for pemphigoid disease.

### Successful inhibition of BTK extends to K/BxN STA and ITP

To assess whether the positive effects of BTK inhibition in the different EBA models extend to other immune complex-mediated autoimmune diseases, ibrutinib was tested in K/BxN STA and ITP. K/BxN STA was induced by the injection of K/BxN serum and ibrutinib was administered daily. The mice were observed for seven days (Fig. 6a). The arthritis score stagnated for the treated mice after about three days, and a significant difference compared to the control mice was observed (Fig. 6b). The levels of inflammatory cytokines in the serum were assessed, and three cytokines were significantly increased on day 7 in treated mice, IL-12p70, IL-23, and IL-27 (Fig. 6c). On day 0, the only difference was observed for IL-1α, which was decreased in the treated mice (Table 3). However, biological significance of these differences is likely minor due to the low overall cytokine concentrations observed in this model. The mitigated disease is also represented by clinical images of the paws, by H&E staining showing the influx of immune cells to the site of inflammation and Safranin-O staining demonstrating the proteoglycan degradation in the control mice compared to the ibrutinib-treated animals (Fig. 6d).

**Fig. 6:**
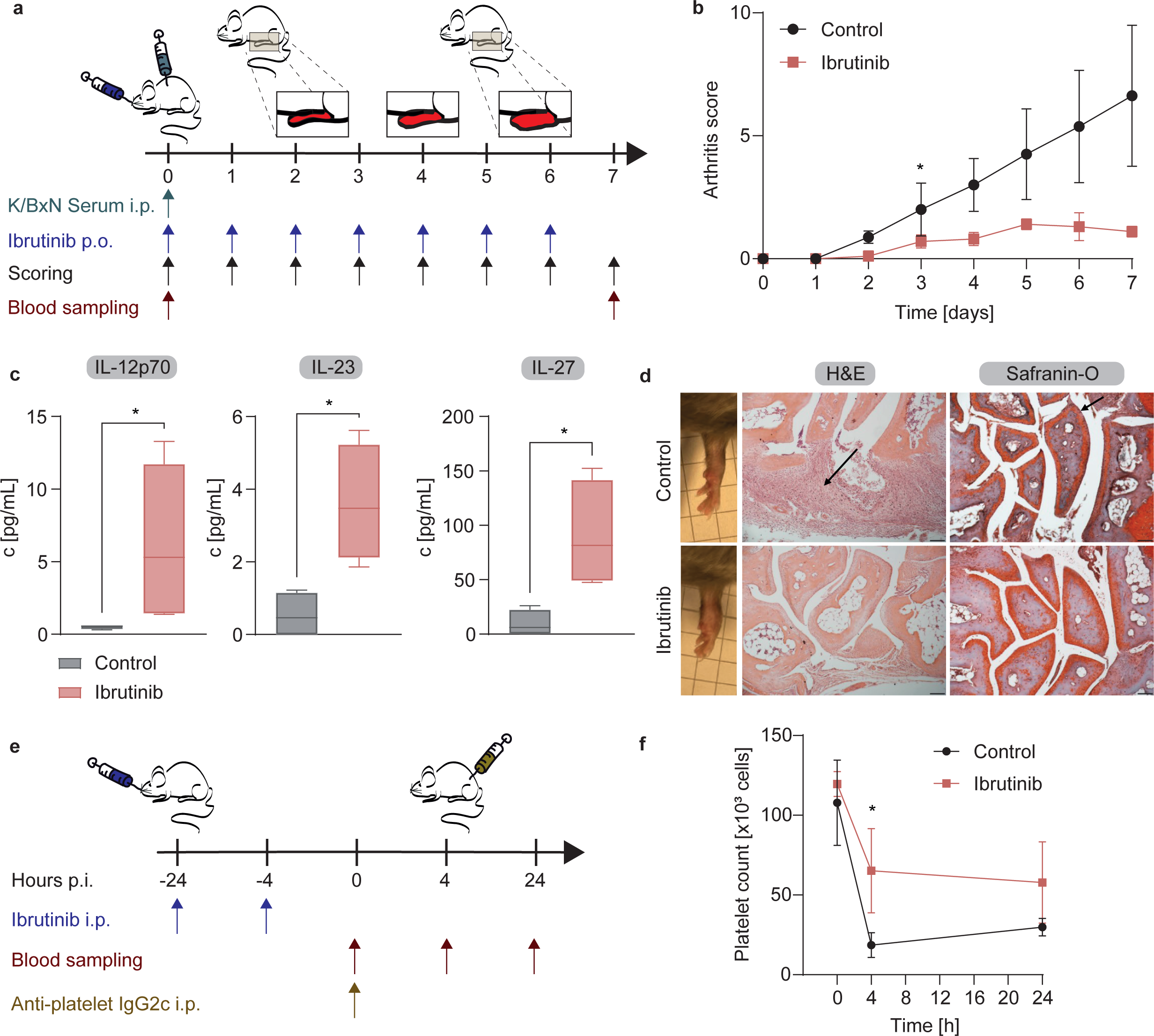
Ibrutinib improves both K/BxN STA and ITP. **(a)** Mice were treated with K/BxN serum to induce STA and injected daily with ibrutinib. **(b)** The swollen paws were evaluated, and the arthritis score was determined. Data are represented as mean ± SD. **(c)** Serum from day 7 was analyzed for cytokine concentrations. Individual values and mean are depicted. **(d)** Representative clinical and histological images are depicted. Arrows indicate cell infiltration (H&E) and proteoglycan degradation (Safranin-O). **(e)** For ITP, mice were pre-treated with ibrutinib, and the disease was induced by anti-platelet IgG2c. **(f)** The platelet count was determined. Data are represented as mean ± SD. * p ≤ 0.05. n = 5 (ibrutinib), n = 4 (control).

In ITP, treatment with ibrutinib was initiated 24 h before disease induction with anti-platelet IgG2c (Fig. 6e). The platelet depletion was significantly inhibited by ibrutinib after 4 h. After 24 h, there was no difference between the groups regarding platelet numbers, but a trend towards higher platelet counts in the treated mice (Fig. 6f). These two additional models demonstrate the power of BTK inhibition in immune complex-mediated autoimmune diseases.

## Discussion

Signal transduction is a major cellular process that determines how a cell responds to a specific stimulus. In IC-mediated autoimmune diseases, FcγRs on the surface of immune cells such as neutrophils are activated, instigating downstream kinase signaling pathways, which in turn result in tissue damage. While most studies focus on investigating the response to non-specific stimuli such as phorbol 12-myristate 13-acetate or N-formylmethionyl-leucyl-phenylalanine ^37–40^, the present study aims to uncover more physiologic FcγR-mediated signaling. To the best of our knowledge, we are the first to perform an unbiased library screening combined with a kinase activity profiling instead of a candidate gene approach. We identified 51 modulated kinases, 44 of which had not previously been described in this context, and 18 targets were functionally validated. Putative targets were further evaluated in preclinical model systems, including therapeutic application, as well as human immune system mice. Thus, we confirmed molecules previously linked to FcγR signaling, including p38 and BTK, and identified PLC, PDK-1, PKC, DNA-PK, KSP, c-Met, and TBK-1 as additional potential therapeutic targets. Furthermore, BTK emerged as an important player validated across disease models. Thus, we contributed to the detailed investigation of FcγR-mediated signaling in neutrophils.

Importantly, human neutrophils have been shown to express different FcγRs ^41^. While signaling may differ for each individual receptor, the interplay between the individual receptors determines the final outcome. Upon crosslinking of activating FcγRs in mice and humans, except for the human glycosylphosphatidylinositol-anchored FcγRIIIB, immunoreceptor tyrosine-based activation motifs for activating (ITAMs) are phosphorylated by SRC family kinases ^42^. Subsequently, SYK is activated, which has previously been shown to play a crucial role in EBA ^34,43^, ITP ^44^ and RA^45^ pathogenesis. SYK can then trigger different interconnected signaling pathways, for example the PI3K/AKT/mTOR, PLC, ERK, and p38 pathways ^42^. These signaling networks determine the cellular activity, such as cytokine release, cytoskeletal changes, transcriptional activity, or cell death ^43^. In this study, we demonstrate that further signaling molecules, including BTK, PLC, KSP, DNA-PK, PKC, PDK-1, p38, and TBK-1 are involved in IC-induced neutrophil signaling (Fig. 7). Additionally, c-Met, also known as hepatocyte growth factor (HGF) receptor, was effectively inhibited. This membrane-bound receptor tyrosine kinase can activate SRC ^46^, which in turn leads to FcγR activation ^41^. In addition, c-Met can also trigger FcγR-independent signaling by direct activation of the PI3K/AKT pathway, FAK, JNK, Ras, PKC ^46^. Human polymorphonuclear granulocytes (PMNs) store HGF in their secretory vesicles and gelatinase/specific granules which is released upon degranulation ^47^. Therefore, this is an alternative pathway triggering neutrophil effector functions upon previous activation.

**Fig. 7:**
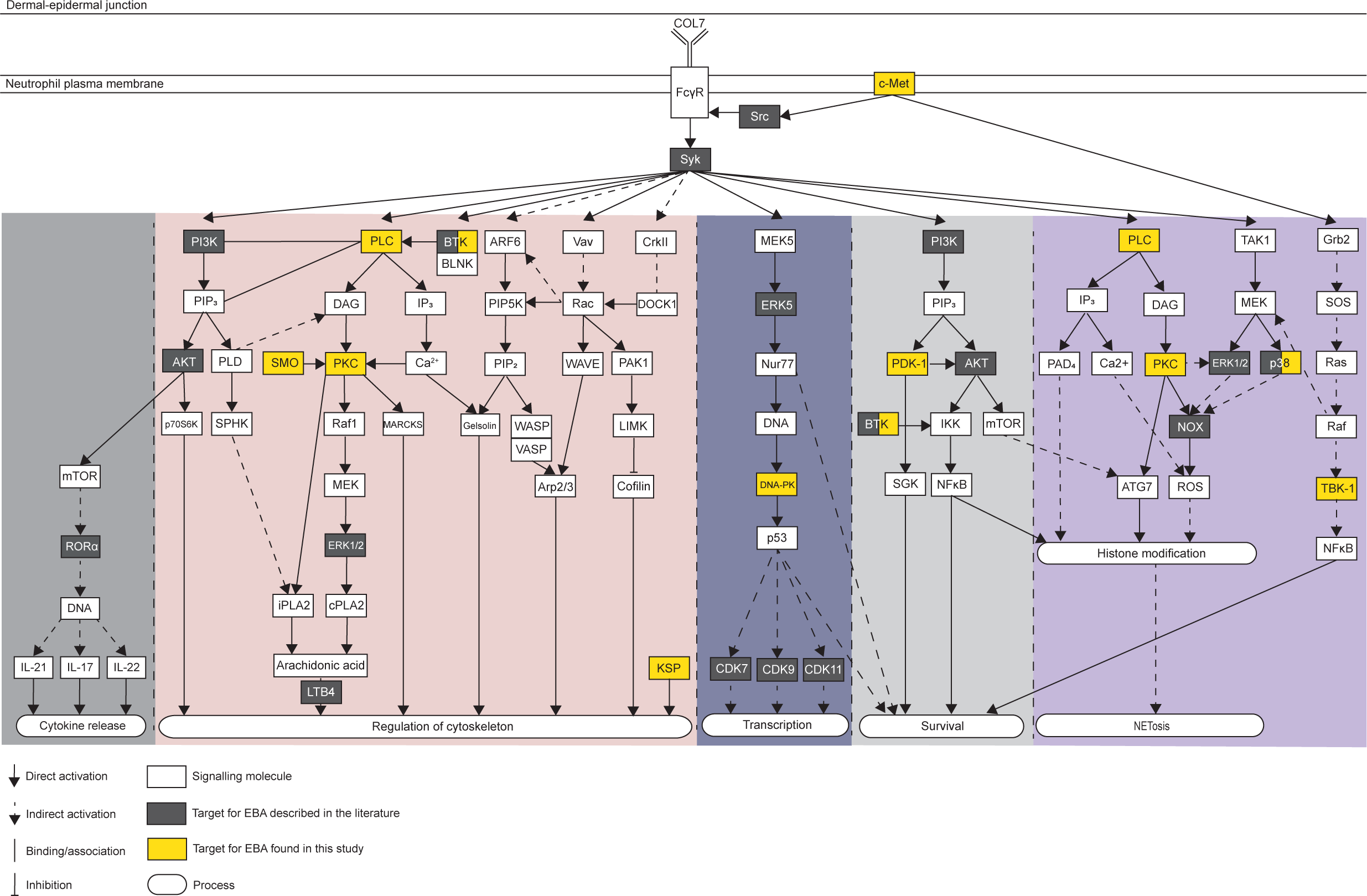
Immune complex-induced signal transduction in neutrophils. Signaling cascades upon immune complex-induced FcγR activation in neutrophils are shown. Pathway data were obtained from the KEGG database (maps 04613, 04666, and 04210) and published literature, and supplemented with findings from this study. The scheme represents our current understanding and may not be exhaustive.

Based on the screening of an STI library and a multiplex kinase activity profiling using IC-stimulated human neutrophils, we identified potential novel therapeutic targets for neutrophil-mediated IC-induced autoimmune diseases. In addition, targets that have already been demonstrated to be implicated in EBA signaling were confirmed including PI3K ^30–32^, AKT1 ^26^, SYK ^33,34^, p38 ^26^, ERK1/2 ^26^, BTK ^27^, CDKs ^28^ and JAK2 ^48^, further strengthening the validity of our results. As stated above, SYK inhibition was also effective in a model of RA^45^ and is approved as treatment in ITP ^44^. Furthermore, we have previously demonstrated the efficacy of CDK9 inhibition in a model of RA ^28^. Of note, the ERK5 inhibitor XMD8-92, which is described to improve experimental EBA ^29^, marginally exceeded the threshold in the library screen by 1.7 %. This may be due to the small sample size during screening and therefore supports our findings. These initial analyses were complemented by an RNAseq analysis revealing changes in the targets also on gene expression level. Several inhibitors (U73122, BX-795, sotrastaurin, BIRB 796, NU7441, SB 743921 HCl, PHA-665752, ibrutinib, and GSK8612) were successfully validated both *in vitro* and *in vivo* in models of EBA.

All of these inhibitors reduced IC-mediated neutrophil activity *in vitro*. Importantly, IL-8-induced chemotaxis was additionally reduced by several STIs, suggesting that FcγR-independent pathways are also at least partially involved in IC-induced autoimmune diseases such as EBA and could contribute to therapeutic efficacy. However, significant effects were only observed at the highest inhibitor concentration except for the PLC inhibitor U73122. The consistent effects of several BTK inhibitors further strengthens the mechanistic interpretation.

Since the BTK inhibitor ibrutinib was the most effective out of the tested STIs, it was applied in further models of IC-mediated autoimmune diseases. Consistent with an independent report ^27^, ibrutinib markedly reduced disease manifestation in local EBA in a prophylactic approach. Notably, we demonstrate for the first time that ibrutinib impaired immunization-induced EBA in a therapeutic setting, more closely reflecting the human disease. Given that this model also involves the immune cells relevant to the afferent phase of EBA, it is complementary to the antibody transfer-induced model, further supporting BTK inhibition as a promising novel treatment option in patients. The translational potential of ibrutinib is underscored by its efficacy in the HIS model where disease development depends on human immune cells and FcγRs. Previously, this model has also been used to demonstrate the therapeutic efficacy of the PDK-1 inhibitor BX-795, further validating our results ^36^.

To explore the role of BTK beyond EBA, ibrutinib was applied in IC-mediated autoimmune disease models of RA and ITP, where it also effectively reduced disease severity. This reduction underlines the importance of BTK in IC-dependent diseases. However, the decrease in classical monocyte numbers in the skin of HIS mice and the efficacy of ibrutinib in macrophage-mediated ITP suggests that the effects are not limited to neutrophils. Consistent with these results, other BTK inhibitors are already undergoing clinical trials for ITP ^49^. Given that ibrutinib potently reduced disease severity in all models, this suggests a consistent mechanism of action across IC-mediated diseases.

Despite the successful application of nine STIs in local EBA as systemic treatment, six inhibitors that were effective in the *in vitro* models did not translate into the *in vivo* models. This may be explained by

at least four causes: first, the species difference of the cells. Human neutrophils were utilized *in vitro* due to better availability and to mimic the patient situation. Despite these advantages, FcγRs and IgG subclasses, among others, differ between humans and mice possibly accounting for discrepancies between the models. Second, only neutrophils were analyzed *in vitro*, but several additional cell types, like eosinophils and lymphocytes ^18^, are involved in PD pathogenesis. While studying neutrophils as a surrogate can be a good model to predict therapeutic efficacy *in vivo* ^30,34,50^, the effect of their interplay with other cell types is neglected. Of note, distinct FcγRs are expressed to different extents on the individual immune cell types. In humans, neutrophils mainly express the signaling-deficient receptors FcγRIIIb and activating FcγRIIA, whereas activating FcγRI and the inhibitory FcγRIIb are present at low levels ^41,51^. In mice, a unique characteristic of neutrophils is their exclusive expression of FcγRIIIb, primarily alongside FcγRIV and the inhibitory FcγRIIb. In contrast, murine monocytes express FcγRI, FcγRIII and FcγRIIb with additional expression of FcγRIV on non-classical monocytes, while human monocytes prominently express FcγRIIa and FcγRI (classical monocytes) or FcγRIIIa (non-classical monocytes) ^42^, suggesting that activation of these receptors triggers different signaling events resulting in heterogeneous outcomes. Third, neutrophils were stimulated *in vitro* with immobilized ICs as those are present in EBA. In addition, soluble ICs also play a role in RA. In contrast, antibodies directly bind to thrombocytes in ITP, causing a potential difference in FcγR signaling via distinct receptor crosslinking ^52^. Fourth, the bioavailability of the inhibitors might not be ideal, preventing a therapeutic effect even if inhibition of the target was complete in an *in vitro* setting.

Another limitation is the use of STIs to study signal transduction, since no inhibitor is perfectly selective and off-target activity cannot be excluded. For most targets, different STIs with distinct selectivity profiles are available, allowing for further exploration in the future. Another option is proteolysis-targeting chimeras (PROTACs) which lead to the specific proteasome-dependent degradation of the target ^53^. This option leaves room for future exploration, provided that more PROTACs become available. In addition, published concentrations with known efficacy were used for the *in vivo* experiments, reducing the risk for adverse events, but not allowing to find the optimal dose. An alternative to STIs for research purposes would be genetic ablation, yet this often results in embryonic lethality and induces a severe burden in the animals ^54,55^. In contrast, STIs are a clinically relevant option, especially for topical applications in dermatological diseases such as pemphigoid.

Together, the present work advances our understanding of IC-induced neutrophil signaling and identifies selective kinase inhibition, especially BTK inhibition, as a promising therapeutic avenue for IC-mediated neutrophil-dependent autoimmune diseases. By providing new mechanistic insights, these findings lay the foundation for future translational efforts aimed at developing more targeted and effective therapeutic strategies.

## Methods

### Sampling of human biomaterials

Whole blood was obtained from healthy volunteers following the provision of written informed consent. All experiments using human samples were approved by the local ethics committee (AZ #20-338, University of Lübeck, Lübeck, Germany) and were performed in accordance with the Declaration of Helsinki.

### Chemicals

All standard chemicals were supplied by either Carl Roth (Karlsruhe, Germany) or Sigma-Aldrich (Taufkirchen, Germany). STIs were supplied by Selleck Chemicals (Houston, Texas, USA) or MedChemExpress (Monmouth Junction, New Jersey, USA). For the initial STI screening, the L3500 target-selective inhibitor library (Selleck Chemicals) was used. Additionally, a curated subset of STIs was selected (Supplementary table 1). For *in vitro* assays, STIs were dissolved at a concentration of 10 mM in dimethyl sulfoxide (DMSO), except for U73122, which was dissolved in dimethylformamide (DMF), and then diluted to achieve final concentrations of 10 µM, 1 µM, 0.1 µM, and 0.01 µM. If further concentrations were required to achieve dose dependency, lower concentrations were included as indicated.

### Analysis of RNAseq data

Data for RNAseq analysis were retrieved from the European Nucleotide Archive (ENA) at EMBL-EBI under accession number PRJEB103987 (https://www.ebi.ac.uk/ena/browser/view/PRJEB103987) and analyzed as described in the original publication ^28^. Briefly, 100,000 human neutrophils were stimulated with immobilized IC consisting of recombinant human collagen VII (COL7) E-F and anti-human COL7 IgG1 for 6 h at 37 °C and 5 % CO2 as previously described, ^30^ then lysed with Trizol (Invitrogen, Waltham, Massachusetts, USA) and subjected to preparation for RNAseq analysis. Data were analyzed using RStudio (Posit Software, Boston, Massachusetts, USA; version 2024.12.1) as previously described ^50^. Differentially expressed genes were identified using DESeq2 ^56^ on transcripts where the sum of the counts over all samples was > 10, and padj < 0.1 was considered significant.

### PMN purification

For all *in vitro* experiments except the chemotaxis assay, PMNs were isolated from EDTA-treated human whole blood performing a PolymorphPrep™ (Serumwerk Bernburg AG, Bernburg, Germany) density gradient centrifugation as previously described ^30,50^.

For the chemotaxis assay, PMNs were isolated from citrate-treated human whole blood by performing a Ficoll density gradient centrifugation as described before ^30,50^. After 1:2 dilution with 1 % (w/v) polyvinyl alcohol and sedimentation for 25 min, the supernatant (up to 40 mL) was transferred on top of a layer of 8 mL of Ficoll Paque PLUS (Cytiva, Marlborough, Massachusetts, USA). Density gradient centrifugation was performed for 24 min at 850 g and room temperature (RT) without brake. For erythrocyte lysis, the granulocyte pellet was resuspended in distilled water, mixed for 45 s, and then neutralized with 2x phosphate-buffered saline (PBS). Two washing steps with PBS were performed for 10 min at 300 g and RT.

The PMNs were diluted to 4×10^6^ cells/mL Neutrophil purity was assessed by flow cytometry for all experiments and was higher than 90 % for all samples.

### Multiplex kinase activity profiling

Multiplex kinase activity profiling was performed as described before ^30^ with modifications using the PamChip® 4 protein tyrosine kinase (PTK) and serine/threonine kinase (STK) peptide microarray system from PamGene International B.V. (BJ‘s-Hertogenbosch, The Netherlands). Human PMNs were purified as described above and diluted in RPMI 1640 without phenol red containing 1 % FCS, 25 mM HEPES, and 2 g/L glucose. Immune complex-coated 6-well plates were prepared as described at a final concentration of 10 µg/mL human COL7 E-F and 2 µg/mL anti-human COL7 IgG1. Per well, 10^7^ cells were added and stimulated for 0 min, 1 min, 5 min, or 15 min at 37 °C and 5 % CO2. Immediately after stimulation the plates were placed on ice. The cell suspension was removed, each well was rinsed with 1 mL cold PBS, and the rinse was added to the cell suspension for each well. The cell suspensions were centrifuged for 5 min at 1000 g and 4 °C. Subsequently, the supernatant was removed, and the pellet was lysed in 100 µL M-PER (Thermo Fisher, Waltham, Massachusetts, USA) for 15 min at RT. The samples were centrifuged for 15 min at 16000 g and 4 °C. Protein concentrations were determined using the Pierce™ BCA Protein Assay Kit (Thermo Fisher) according to the manufacturer’s instructions. Lysates with a concentration > 0.5 µg/µL were included for analysis.

STK and PTK microarray assays were performed according to the manufacturer’s instructions. The activity of kinase was considered to be significantly modulated if the mean specificity score (= negative decadic logarithm of the likelihood of obtaining a higher difference between the groups when assigning peptides to kinases randomly) was > 1 (p < 0.1) and the significance score (= likelihood of obtaining a higher difference for random assignment of values to treatment-and control groups) was > 0.5 (p < 0.32). Unstimulated cells (0 min) were used for comparison with the different time points. The mean kinase statistic was calculated by averaging the difference between the signal intensity of a sample and its control value, normalized against a pooled estimate of the standard deviation in each sample, for each peptide assigned to a specific kinase. Analysis was performed using Bionavigator version 6.3 with the UKA 2018 version (PamGene).

### In vitro assays

For ROS release assay, adhesion assay, and flow cytometry, white flat-bottom high-binding 96-well plates were coated with IC consisting of human COL7 E-F at 2.5 µg/mL and anti-human COL7 IgG1 at 1.8 µg/mL as previously described ^30,57^. PMN were isolated as described above and 2×10^5^ cells per well were added in the presence or absence of STI. Only antibody (with cells), or only antigen (with cells) served as negative controls, while IC (with cells, but without STI) served as positive control. DMSO (or DMF for U73122) was included in all controls at a concentration corresponding to that of the highest inhibitor concentration (0.05 %). All samples were measured in technical duplicates.

### ROS release assay

Following plate preparation as described above, luminol was automatically injected into each well to achieve a final concentration of 90 µg/mL, and chemiluminescence was measured for 2 h every 2 min at 37 °C in a GloMax® Discover microplate reader (Promega, Walldorf, Germany). ROS production was quantified by calculating the AUC, and the results were normalized to the positive control.

### Adhesion assay

Following plate preparation as described above, PMN adhesion was assessed using an impedance-based real-time cell analysis assay, as previously described ^58,59^. Briefly, a 96-well E-plate (Agilent, Santa Clara, California, USA) was coated with IC as described for the ROS release assay. PMN adhesion was monitored using the xCELLigence Real-Time Cell analysis system (Agilent), with impedance measurements recorded every 2 min for 120 min. Adhesion was expressed as the cell index, an arbitrary unit reflecting changes in electrical impedance. Data were analyzed by calculating the AUC, and the results were normalized to the positive control, as described for the ROS release assay.

### Flow cytometry

Following plate preparation as described above, human PMNs were stimulated with IC for 2 h at 37 °C and 5 % CO₂. Subsequently, they were stained for the flow cytometric analysis of the following surface markers using standard flow cytometry procedures: CD14 (clone HCD14), CD16 (clone 3G8), CD18 (clone 1B4/CD18), CD45 (clone HI30), CD62L (clone DREG-56), and CD193 (clone 5E8). Viability was assessed by Annexin V (B266195) and Zombie NIR staining ^50^. All dyes were purchased from BioLegend (San Diego, California, USA). Analyzes for activation markers were performed on FSC-SSC gated CD45^+^ singlets. Annexin V^-^Zombie NIR^-^cells were considered viable. Measurements were performed on a MACSQuant® Analyzer 10 (Miltenyi Biotec, Bergisch Gladbach, Germany), and analysis was conducted using MACS Quantify software (version 2.13.3, Miltenyi Biotec).

### Trypan blue staining

To test the toxicity of compounds which interfered with the flow cytometric assay due to their intrinsic color, a trypan blue staining was performed for those. Plate preparation, neutrophil isolation, and incubation were conducted as for the flow cytometric approach. Following incubation, the cell suspensions were stained using trypan blue at a final concentration of 0.04 % (v/v) and the percentage of alive cells was microscopically assessed.

### Chemotaxis

Human PMNs were analyzed for their IL-8-induced chemotactic ability using a 48-well Boyden chamber as described before ^50^. Briefly, IL-8 (Thermo Fisher) was used as stimulus at 5-15 nM depending on the batch, so that at least 30 % migration was observed in the positive control. STIs were prepared in 1 (w/v) % BSA in PBS and mixed 1:1 with the cell suspension. The STI-containing cell suspension was applied to the upper chamber at 50 µL per well and incubated for 1 h at 37 °C in a humid chamber. Buffer only and IL-8 only with cells, but without STIs served as negative and positive control, respectively. Normalization was conducted to the positive control (IL-8 with cells only) for each individual experiment.

### Animal study approval

Animal experiments were approved by local authorities of the Animal Care and Use Committee (government of Schleswig-Holstein and government of Lower Franconia) and the Institutional Animal Care and Use Committee at Rush University. All experiments were performed by certified personnel (AZ 108/08-15, AZ 97-11/20, 55.2.2-2532-2-2085, and 55.2.2-2532-2-2146) following the ARRIVE guidelines.

### Animal experimentation

Mice were bred in a specific pathogen–free environment and provided standard mouse chow and chlorinated water *ad libitum*. Clinical examinations performed under anesthesia, using intraperitoneal (i.p.) administration of a ketamine (75 mg/kg, Sigma-Aldrich) and medetomidine (1 mg/kg, Vetoquinol, Ismaning, Germany) mixture, antagonized after ca. 30 min by atipamezole (5 mg/kg, Vetoquinol) or isoflurane. On the final day, the mice were anesthetized using 15 mg/kg xylazine (Sigma-Aldrich) and 100 mg/kg ketamine and killed by cervical dislocation or CO2.

### Local antibody-transfer induced experimental EBA

Specific anti–mouse COL7^C^ IgG from rabbit serum was isolated as previously described ^12,60^. Experimental EBA was induced in C57BL/6J mice (Jackson Laboratory, Bar Harbor, Maine, USA) of both sexes aged 8 to 14 weeks by ear-base injection with 100 μg rabbit anti–mouse COL7^C^ IgG. For systemic treatment, STIs were administered as indicated in Table 1 at 100 µL/20 g mouse. Concentrations were adapted from established published protocols (Supplementary table 1). For topical treatment in independent experiments, STIs were applied as indicated in Table 2, and treatment was performed daily for four days. Respective vehicle-treated mice served as controls in all cases. The mice were clinically evaluated every day during the five-day experiment, and the percentage of ear thickness was assessed using a Mitutoyo 7301 dial thickness gauge (Neuss, Germany), as well as the AESA, all by a blinded person. Ears were collected on the final day of the experiment, and half of each ear was fixed in 4 % histofix and embedded paraffin according to standard protocols or frozen in liquid nitrogen and stored at -80 °C. Histological analysis of H&E staining and direct immunofluorescence staining for murine IgG and C3 were performed as described elsewhere ^12^. Scoring of H&E-stained sections was performed by blinded observers to determine (i) the magnitude of epidermal thickness with values from 0 (10-20 µm), 1 (20-40 µm), 2 (40-100 µm) to 3 (> 100 µm), (ii) the percentage of dermal-epidermal split formation (length of split formation/length of dermal-epidermal junction), with values from 0 (no split), 1 (< 20 %), 2 (20-50 %) to 3 (> 50 %) and (iii) the dermal infiltration with values from 0 to 3 corresponding to no, mild, moderate or severe degree, respectively. The H&E score was calculated as the mean of these three parameters. For all clinical and histological parameters assessed, the mean of both groups was calculated and the percentage of reduction in the treatment group compared to the control group was determined.

**Table 1.**
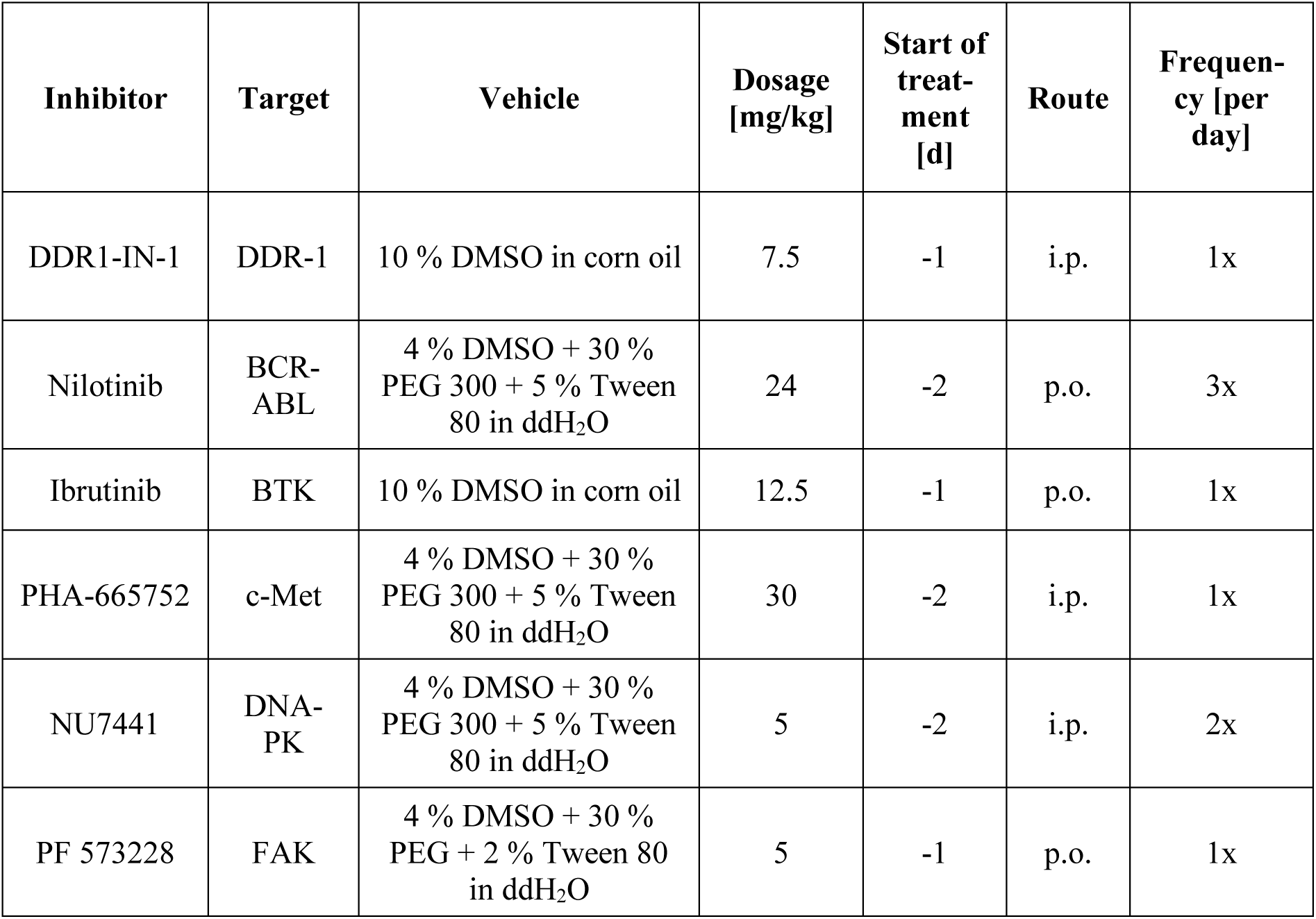
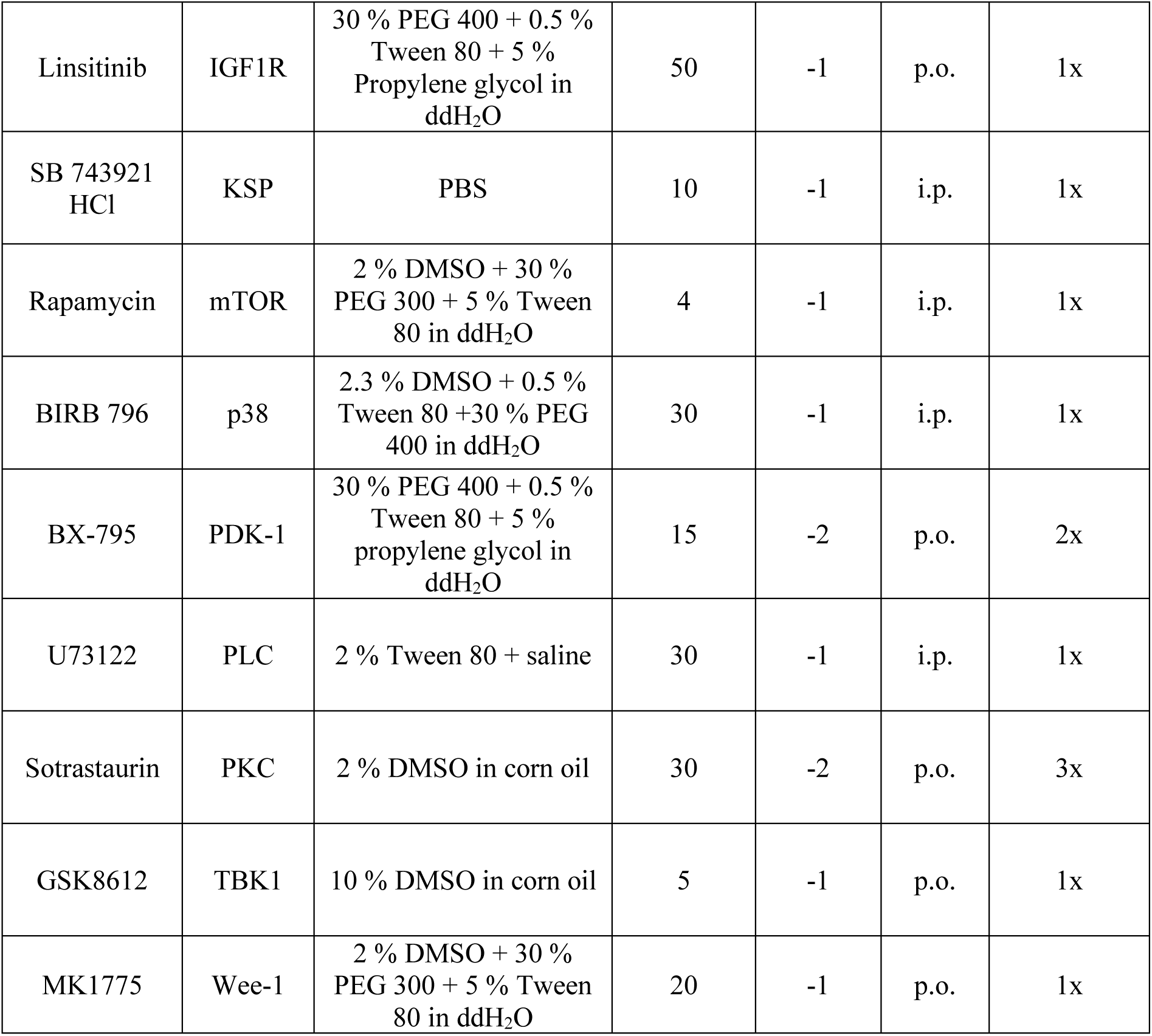
Systemic application of STIs in local antibody transfer-induced EBA. The target, vehicle, dosage, treatment schedule, and route of administration are indicated for each STI. i.p., intraperitoneally; p.o., *per os*.

| Inhibitor | Target | Vehicle | Dosage [mg/kg] | Start of treatment [d] | Route | Frequency [per day] |
| --- | --- | --- | --- | --- | --- | --- |
| DDR1-IN-1 | DDR-1 | 10 % DMSO in corn oil | 7.5 | -1 | i.p. | 1x |
| Nilotinib | BCR-ABL | 4 % DMSO + 30 % PEG 300 + 5 % Tween 80 in ddH<sub>2</sub>O | 24 | -2 | p.o. | 3x |
| Ibrutinib | BTK | 10 % DMSO in corn oil | 12.5 | -1 | p.o. | 1x |
| PHA-665752 | c-Met | 4 % DMSO + 30 % PEG 300 + 5 % Tween 80 in ddH<sub>2</sub>O | 30 | -2 | i.p. | 1x |
| NU7441 | DNA-PK | 4 % DMSO + 30 % PEG 300 + 5 % Tween 80 in ddH<sub>2</sub>O | 5 | -2 | i.p. | 2x |
| PF 573228 | FAK | 4 % DMSO + 30 % PEG + 2 % Tween 80 in ddH<sub>2</sub>O | 5 | -1 | p.o. | 1x |
| Linsitinib | IGF1R | 30 % PEG 400 + 0.5 %<br>Tween 80 + 5 %<br>Propylene glycol in<br>ddH<sub>2</sub>O | 50 | -1 | p.o. | 1x |
| SB 743921<br>HCl | KSP | PBS | 10 | -1 | i.p. | 1x |
| Rapamycin | mTOR | 2 % DMSO + 30 %<br>PEG 300 + 5 % Tween<br>80 in ddH<sub>2</sub>O | 4 | -1 | i.p. | 1x |
| BIRB 796 | p38 | 2.3 % DMSO + 0.5 %<br>Tween 80 + 30 % PEG<br>400 in ddH<sub>2</sub>O | 30 | -1 | i.p. | 1x |
| BX-795 | PDK-1 | 30 % PEG 400 + 0.5 %<br>Tween 80 + 5 %<br>propylene glycol in<br>ddH<sub>2</sub>O | 15 | -2 | p.o. | 2x |
| U73122 | PLC | 2 % Tween 80 + saline | 30 | -1 | i.p. | 1x |
| Sotrastaurin | PKC | 2 % DMSO in corn oil | 30 | -2 | p.o. | 3x |
| GSK8612 | TBK1 | 10 % DMSO in corn oil | 5 | -1 | p.o. | 1x |
| MK1775 | Wee-1 | 2 % DMSO + 30 %<br>PEG 300 + 5 % Tween<br>80 in ddH<sub>2</sub>O | 20 | -1 | p.o. | 1x |

**Table 2.** Topical application of STIs in local antibody transfer-induced EBA. The target, vehicle, dosage and treatment frequency are stated for each inhibitor. DAC, *Deutscher Arzneimittel-Codex*/German Drug Codex.

| Inhibitor | Target | Vehicle | Dosage | Frequency<br>[per day] |
| --- | --- | --- | --- | --- |
| Ibrutinib | BTK | acetone/DMSO (20:1) | 65 µg/100 µL | 1x |
| PHA-665752 | c-Met | DMSO/DAC base cream (1:1) | 6.4 % | 2x |
| NU7441 | DNA-PK | acetone/DMSO (20:1) | 65 µg/100 µL | 1x |
| SB 743921<br>HCl | KSP | acetone/DMSO (20:1) | 156.25 µg/100 µL | 1x |
| BIRB 796 | p38 | acetone/DMSO (20:1) | 156.25 µg/100 µL | 1x |
| BX-795 | PDK-1 | DMSO/DAC base cream (1:1) | 5 % | 2x |
| U73122 | PLC | acetone/DMSO (20:1) | 156.25 µg/100 µL | 1x |
| Sotrastaurin | PKC | DMSO/DAC base cream (1:1) | 4.35 % | 2x |
| GSK8612 | TBK1 | acetone/DMSO (20:1) | 65 µg/100 µL | 1x |

**Table 3.** Descriptive statistics for all cytokines analysed in K/BxN STA at day 0 and day 7. Mann-Whitney test was performed to compare the treatment and control groups at each time point, and the p values are indicated.

|  | day 0 |  |  | day 7 |  |  |
| --- | --- | --- | --- | --- | --- | --- |
|  | median ± SEM [pg/mL] |  | p value | median ± SEM [pg/mL] |  | p value |
| cyto-<br>kine | treated | control | treated vs.<br>control | treated | control | treated vs.<br>control |
| GM-CSF | 0.4600 ±<br>0.2524 | 0.03500 ±<br>0.03500 | 0.1032 | 1.848 ±<br>0.9854 | 0.09000 ±<br>0.05212 | 0.1032 |
| IL-10 | 38.84 ±<br>17.63 | 13.82 ±<br>3.532 | 0.4127 | 13.14 ±<br>3.745 | 18.04 ±<br>5.319 | 0.6857 |
| IL-12p70 | 1.986 ±<br>0.5247 | 1.213 ±<br>0.3885 | 0.4127 | 6.320 ±<br>2.363 | 0.5025 ±<br>0.07075 | 0.0159 |
| IL-17A | 0.7200 ±<br>0.1381 | 1.075 ±<br>0.1686 | 0.4524 | 0.6060 ±<br>0.7067 | 0.3275 ±<br>0.04385 | 0.1111 |
| IL-1α | 9.642 ±<br>0.4692 | 136.3 ±<br>91.99 | 0.0159 | 4.338 ±<br>0.7106 | 9.053 ±<br>4.611 | 0.7429 |
| IL-1β | 5.544 ±<br>1.099 | 8.803 ±<br>7.940 | 0.2540 | 13.11 ±<br>5.073 | 2.453 ±<br>0.4762 | 0.1667 |
| IL-23 | 6.700 ±<br>4.771 | 0.000 ±<br>0.000 | 0.2857 | 3.608 ±<br>0.8051 | 0.5350 ±<br>0.3149 | 0.0286 |
| IL-27 | 119.5 ±<br>75.86 | 28.81 ±<br>17.14 | 0.3492 | 90.71 ±<br>24.67 | 9.530 ±<br>5.979 | 0.0286 |
| IL-6 | 0.5860 ±<br>0.1182 | 0.2975 ±<br>0.1158 | 0.1270 | 0.8525 ±<br>0.3165 | 4.833 ±<br>2.433 | 0.0857 |
| IFN-β | 3.500 ±<br>2.282 | 1.625 ±<br>0.7028 | 0.7429 | 2.875 ±<br>0.8189 | 5.175 ±<br>2.622 | 0.8857 |
| IFN-γ | 4.450 ±<br>0.6046 | 2.870 ±<br>0.6064 | 0.1667 | 7.072 ±<br>2.301 | 3.835 ±<br>0.6288 | 0.5556 |
| MCP-1 | 2.430 ±<br>1.071 | 2.738 ±<br>1.666 | 0.9048 | 6.073 ±<br>1.612 | 7.443 ±<br>1.041 | 0.5714 |
| TNF-α | 6.498 ±<br>0.5475 | 6.420 ±<br>1.694 | > 0.9999 | 11.13 ±<br>1.398 | 7.000 ±<br>2.430 | 0.2857 |

### Immunization-induced experimental EBA

B6.SJL-H2b C3c/2CyJ mice (B6.s, Jackson Laboratory) mice were immunized with 120 µg of von Willebrand factor A-like domain 2 (vWFA2) of mCOL7 (mCOL7^vWFA^^2^) diluted 1:2 in Titermax (Sigma-Aldrich) as previously described ^12,61^. Mice were randomized into treatment groups once 2 % or more of the body surface area were affected. Ibrutinib was applied at 90 mg/kg body weight twice daily p.o. for four weeks. Blinded clinical scoring was performed once weekly. Serum samples were obtained before immunization, after immunization but before randomization, and after 2 and 4 weeks of treatment to determine circulating total and specific IgG. Clinical scores were normalized to the initial clinical score when allocated into treatment groups. Circulating total and autoreactive anti-mCOL7^vWFA^^2^ IgG in plasma from immunized mice were quantified by ELISA using mouse IgG quantification sets (Bethyl, Montgomery, Texas, USA) according to the manufacturer’s instructions with minor modifications ^62^. Specifically, wells were coated with 250 ng of recombinant mCOL7^vWFA^^2^ in coating buffer. After blocking, diluted samples were incubated for 60 min. Bound IgG was detected with HRP-conjugated goat anti-mouse antibodies (Bethyl) and tetramethylbenzidine (Invitrogen). The reaction was stopped with 2 M sulfuric acid, and absorbance was read on a GloMax® Discover microplate reader (Promega) at 450 nm. Concentrations were calculated from standard curves generated with the provided mouse reference sera (Bethyl). Values were normalized to the timepoint before immunization.

### EBA in HIS mice

A passive antibody transfer model was used to induce an EBA-like phenotype in HIS mice. EBA was induced in HIS mice as described before ^36^. HIS mice were generated as described ^36,63–68^. Briefly, newborn NSG-FcRγ^−/−^ mice were irradiated with a dose of 1.4 Gy and intravenously injected with 20,000–40,000 human hematopoietic stem cells (HSCs) 6-8 h post-irradiation. HSCs were isolated from umbilical cord blood with informed consent from donors as previously described. The efficiency of reconstituting the human immune system was assessed 12 to 14 weeks after transplantation by analysis of human cell subsets in whole blood by flow cytometry as previously described ^36^.

For evaluating the kinase BTK as a therapeutic target, HIS mice were treated orally with ibrutinib (Selleckchem) at 12.5 mg/kg daily. The inhibitor was dissolved in 5 % DMSO, 30 % PEG-300, 0.5 % Tween-80 in water. The buffer solution was also used as vehicle control. Disease progression was monitored by blinded clinical scoring of skin lesions on days 6, 9 and 12 post initial antibody administration. On day 13, mice were sacrificed, and skin samples were collected from the trunk after removing the fur.

Single-cell suspensions for flow cytometry of skin samples were generated by enzymatic digestion as previously described ^36^. Subsequently, the suspensions were processed for flow cytometry analysis. The following antibodies were used for identification of human cells: cFluor V420 anti-CD3 (SK7), cFluor V450 anti-CD14 (M5E2), cFluor V547 anti-human CD45 (HI30), cFluor BYG710 anti-CD19 (HIB19), cFluor R668 anti-CD16 (3G8) and cFluor R720 anti-CD56 (5.1H11) (all from Cytek® cFluor® TBMNK Kit) and BV650 anti-murine CD45 (A20), BV605 anti-CD33 (P67.6) and PE/Cy7 anti-CD66b (G10F5) (BioLegend). For identification of murine immune cells, BV650 anti-murine CD45 (A20), BV421 anti-CD11b (M1/70), Spark Blue 550 anti-Ly6G (1A8), PE-Dazzle594 anti-Ly6C (HK1.4) were used. Cell populations were gated as follows: Aggregates were excluded by forward scatter area/height ratio (FSC-A/FSC-H); dead cells excluded by Zombie NIR staining; human and murine leukocytes were separated with human CD45 against murine CD45. Within the human cells, B cells were identified as CD19^+^ and T cells as CD3^+^; NK cells were gated as CD56^+^ in the CD19^-^CD3^-^population; within the CD56^-^gate, neutrophils were identified as CD66b^+^ and remaining myeloid cells were CD33^+^. Classical monocytes (CD14^+^CD16^-^) and non-classical monocytes (CD14^-^CD16^+^) were gated within the CD33^+^ population. Murine myeloid cells were identified as CD11b^+^. Murine neutrophils were gated as Ly6G^+^ and classical monocytes as Ly6C^+^. Ly6G^-^Ly6C^-^cells were identified as non-classical monocytes. Data was acquired on a Northern Lights flow cytometer (Cytek) and analyzed with FlowJo V10.

Blood samples were obtained on days 0, 3, 6, 9 and 13. Plasma was isolated by centrifugation and stored at -80 °C until cytokine profiling was performed using the LEGENDPlex^TM^ Human Inflammation Panel 1 and Mouse Inflammation Panel to quantify human and murine cytokines, respectively. The data were acquired on the Northern Lights flow cytometer (Cytek) and analyzed with Qognit software to determine cytokine concentrations relative to standard curves.

### K/BxN STA

K/BxN STA was induced by injection of arthritogenic K/BxN serum in female C57BL/6J mice aged 12 weeks at 10 µL/g body weight as described before ^28,29,69^. Before and after arthritis induction, mice were treated daily with 20 mg/kg body weight ibrutinib or vehicle control i.p. The severity of joint swelling was monitored to evaluate clinical disease by a blinded person over the course of seven days. Peripheral blood was collected retro-orbitally on day 0 and day 7, and inflammatory cytokines were quantified in the serum by LegendPlex multiplex assay (Mouse Inflammation 13-plex, BioLegend) as a parameter of systemic inflammation. On day 7, mice were sacrificed, and forelegs were prepared for histological analysis. H&E staining was performed to evaluate inflammatory infiltrates and Safranin-O staining to assess bone and cartilage damage, as previously described ^69,70^. Stained sections were evaluated by light microscopy using a Zeiss Axiovert 200 microscope and the AxioVision LE4.6 software (Olympus Life Science, Waltham, Massachusetts, USA).

### ITP

Platelet depletion in the presence of kinase inhibition was assessed in female C57BL/6J mice. Platelet depletion was induced by injection of 0.35 µg/g anti-platelet IgG2c (clone 6A6) as described ^25,28,29^. Treatment with vehicle control or ibrutinib (12.5 mg/kg body weight p.o.) was performed 24 h and 4 h before the induction of platelet depletion. Platelet counts in peripheral blood were assessed immediately before 6A6-IgG2c injection, and at 4 h and 24 h post-injection in peripheral blood collected retro-orbitally. Platelet counts were determined using an ADVIA® 2120i hematocytometer (Siemens Healthineers, Erlangen, Germany).

### Statistical analysis

Data were analyzed using Prism 10.4.1 (GraphPad Software, San Diego, USA). For all *in vitro* assays, Kruskal-Wallis test with Dunn’s multiple comparisons test was performed for each individual inhibitor and experiment if not indicated differently. Dose response curves and IC50 values were calculated using the non-linear fit inhibitor vs. response (three parameters) function or for EC50 values using the agonist vs. response (three parameters) function where an activating response was observed. For all *in vivo* experiments, two-way analysis of variance (ANOVA) or mixed-effects analysis (in the case of missing values) with Šidák’s multiple comparisons test was performed unless stated otherwise. A value of p ≤ 0.05 was considered significant. The data are displayed as Tukey box-and-whisker plots or else as mean ± standard deviation (SD) unless described differently.

## List of abbreviations

ABSA: affected body surface area
AESA: affected ear surface area
ANOVA: analysis of variance
AUC: area under the curve
B6.s mice: B6.SJL-H2b C3c/2CyJ mice
BSA: bovine serum albumin
COL7: collagen VII
DAC: *Deutscher Arzneimittel-Codex*/German Drug Codex
ddH2O: double-distilled water
DMF: dimethyl formamide
DMSO: dimethyl sulfoxide
EBA: epidermolysis bullosa acquisita
FcγR: fragment crystallizable gamma receptor
H&E: hematoxylin and eosin
HGF: hepatocyte growth factor
HIS: humanized immune system
HSCs: human hematopoietic stem cells
IgG: immunoglobulin G
i.p.: intraperitoneal
IC: immune complex
ITP: immune thrombocytopenia
K/BxN STA: K/BxN serum-transfer arthritis log2
FC: log2 fold change
n.d.: not determined
NET: neutrophil extracellular trap
PBS: phosphate-buffered saline
PD: pemphigoid disease
PMN: polymorphonuclear granulocyte
p.o.: *per os*
PROTAC: proteolysis-targeting chimera
PTK: protein tyrosine kinase
RA: rheumatoid arthritis
RNAseq: bulk RNA sequencing
ROS: reactive oxygen species
RT: room temperature
s.c.: subcutaneously
SD: standard deviation
STK: serine/threonine kinase
STI: signal transduction inhibitor
vWFA2: von Willebrand factor A-like domain 2

## Competing interests

The authors declare that they have no competing interests.

## Funding

This research was funded by the Cluster of Excellence “Precision Medicine in Chronic Inflammation” (EXC 2167), the Research Training Group “Defining and Targeting Autoimmune Pre-Disease” (RTG 2633), and the Collaborative Research Center “Pathomechanisms of Antibody-mediated Autoimmunity” (CRC 1526), all from the Deutsche Forschungsgemeinschaft and the Schleswig-Holstein Excellence-Chair Program from the State of Schleswig Holstein.

## Author’s contributions

Investigation: MS, CO, AK, CZ, MA, GK, KSD, TM, PS, LV, APM, AKS, SMV, YW, HG, JL, writing (original draft): MS, visualization: MS, LV, AKS, formal analysis: MS, CO, AK, NE, AKS, LV, KTA, resources: RV, GV, FP, XY, RJL, KTA, conceptualization: RJL, AL, KB, supervision: AL, KB, KTA, project administration: AL, KB, writing (review & editing): all.

## Acknowledgements

We thank Astrid Fischer, Claudia Kauderer, Alexandra Wobig, Daniela Rieck, Norbert Reiling, Carolin Möller, and Cindy Jensen for their support.

---

**Fig. S1:**
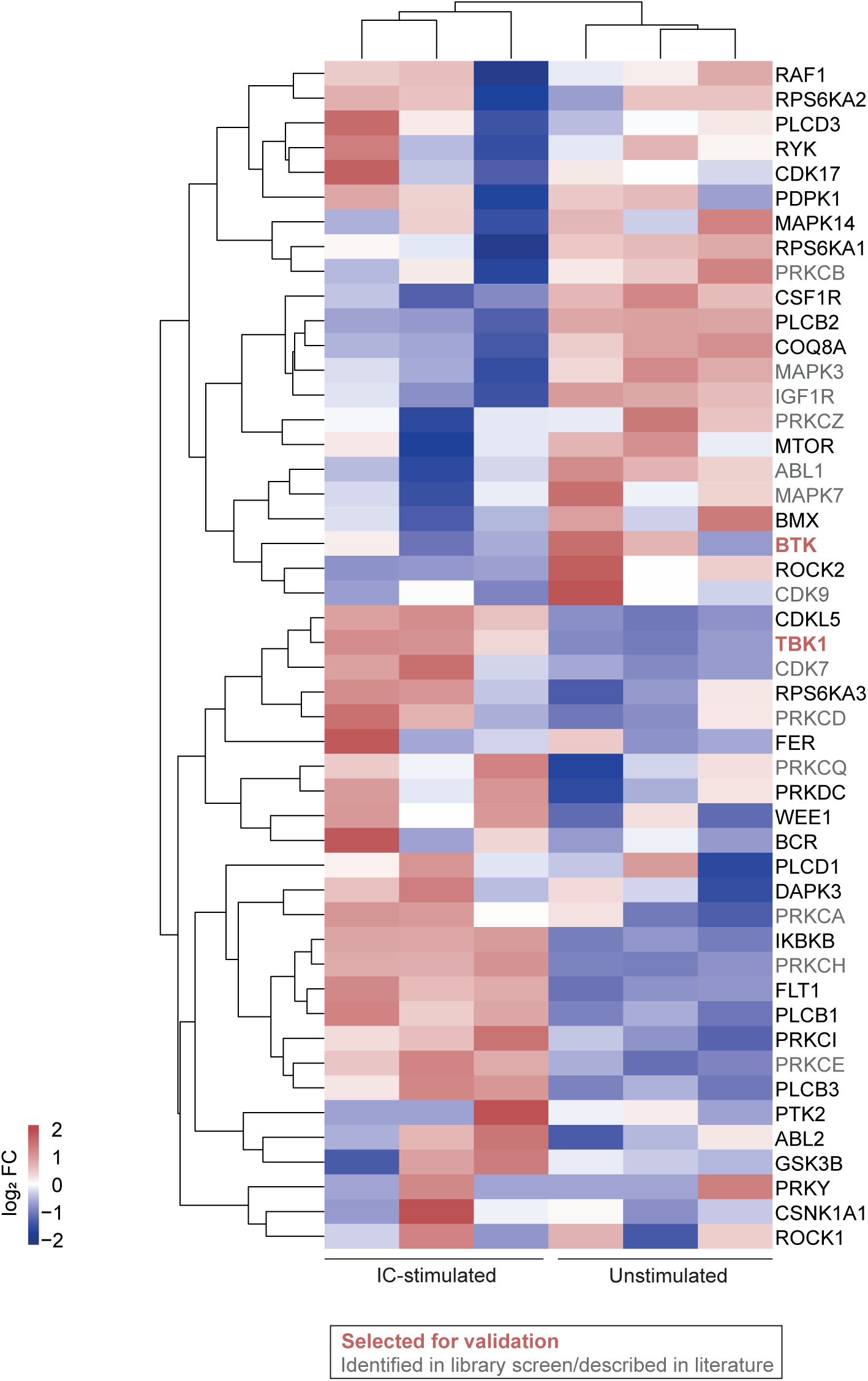
Kinases identified by multiplex kinase activity profiling are differentially expressed at mRNA level upon IC stimulation. Neutrophils isolated from the whole blood of human donors were stimulated for 6 h with immobilized COL7E-F-anti-COL7 IgG1-IC. The cells were lysed and prepared for RNAseq together with unstimulated cells. Differentially expressed genes were identified and log2 fold changes (log2 FC) of kinases identified in multiplex kinases activity profiling are shown.

**Fig. S2:**
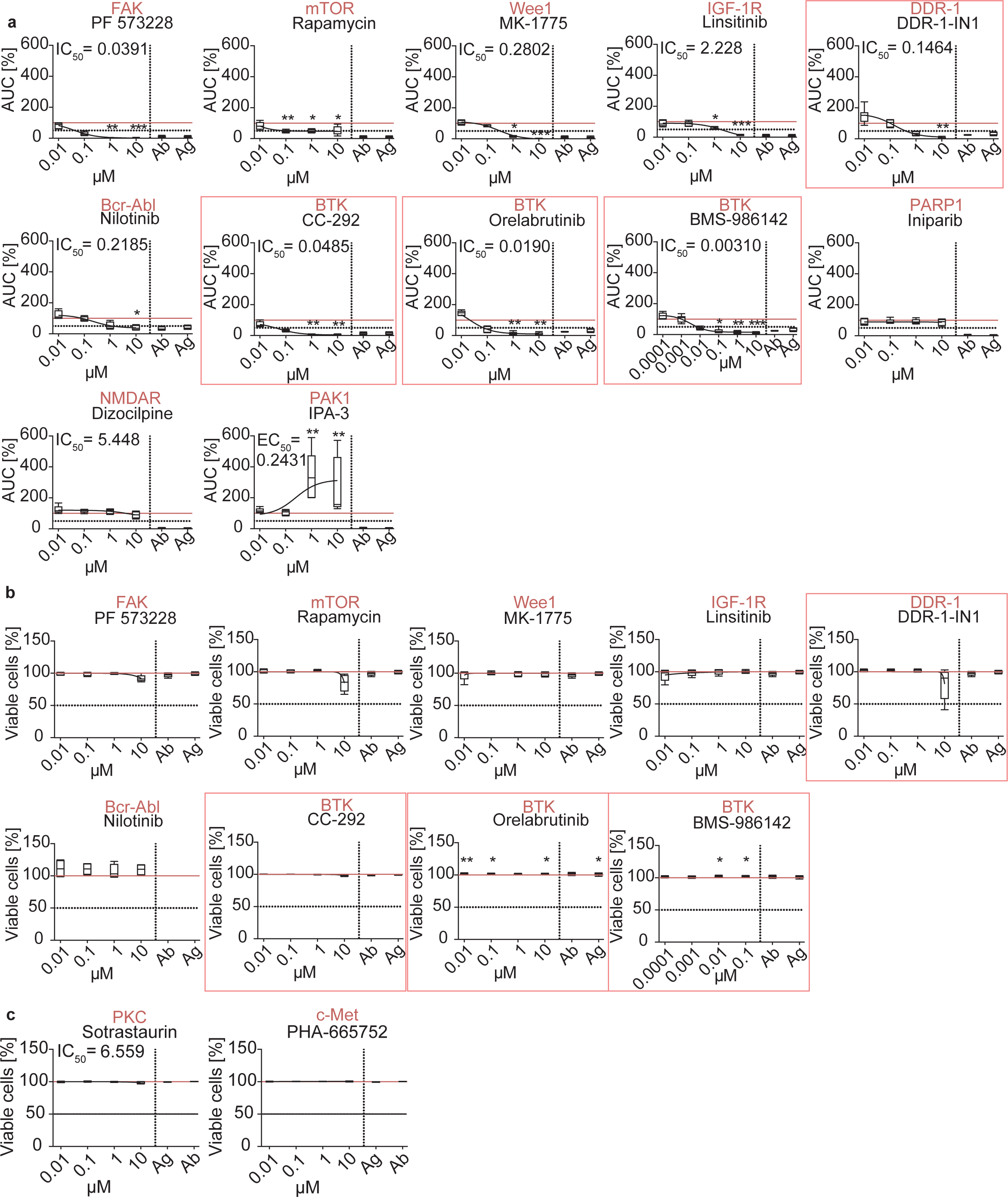
Several additional STIs reduce ROS release from IC-stimulated neutrophils. Human neutrophils treated with immobilized COL7E-F-anti-COL7 IgG1-IC and STIs for 2 h. IC-activated neutrophils treated with DMSO only served as positive control while antibody only (ab)-and antigen only (ag)-treated cells served as negative controls. Normalization was performed to the positive control and is marked by the red horizontal line. STIs that were chosen based on the multiplex kinase activity profiling are highlighted by red boxes. **(a)** ROS release was measured in a luminol-based assay. **(b)** Viability was determined in an Annexin V/Zombie NIR-based assay. **(c)** Viability of colored compounds was assessed by trypan blue staining. n = 5. Kruskal-Wallis test with Dunn’s post-test was performed. * p ≤ 0.05. ** p ≤ 0.01. *** p ≤ 0.001.

**Fig. S3:**
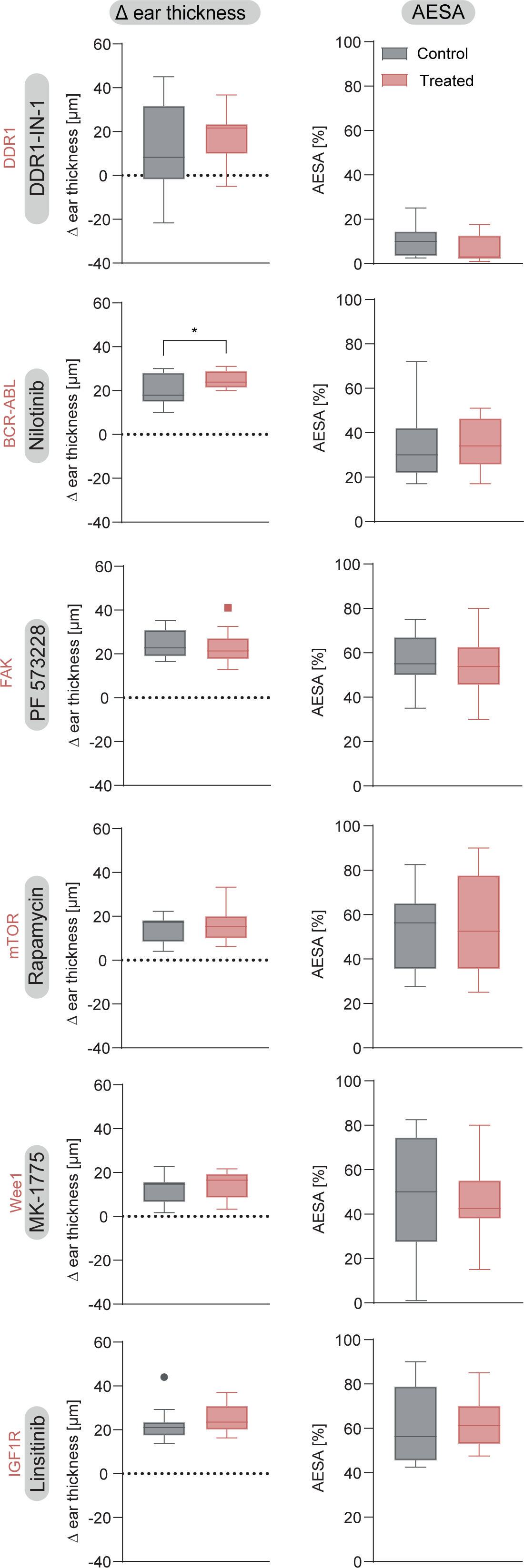
None of the additional STIs tested as systemic treatment in local antibody transfer-induced EBA was effective. Mice were treated with STIs before the induction of local antibody transfer-induced EBA on day 0 and scored daily for five days. The difference in ear thickness and the affected ear surface area (AESA) were determined. Mann-Whitney test was performed. n = 4 (DDR1-IN-1), n = 5 (nilotinib), n = 8 (PF 573228, rapamycin, MK-1775, linsitinib). * p ≤ 0.05. ** p ≤ 0.01. *** p ≤ 0.001.

**Fig. S4:**
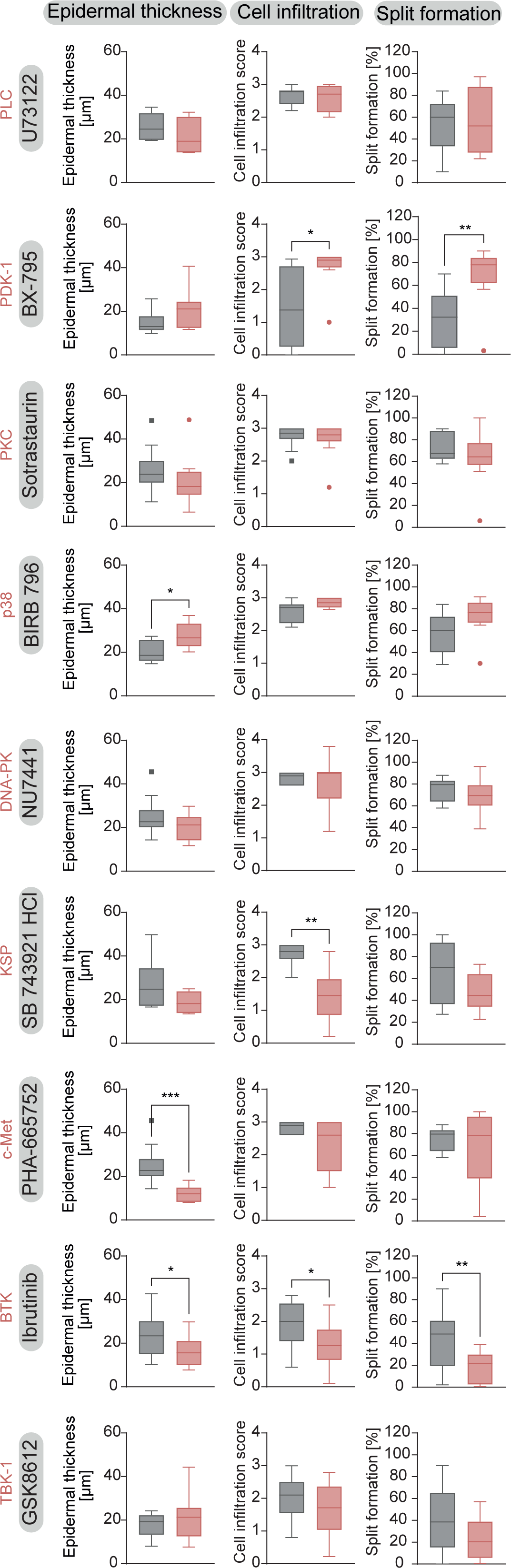
Histological parameters are partially improved by systemic treatment with STIs in local antibody transfer-induced EBA. Mice were treated with STIs before the induction of local antibody transfer-induced EBA on day 0 and clinically scored daily for five days. Epidermal thickness, cell infiltration, and split formation were evaluated using H&E-stained ear sections. * p ≤ 0.05. ** p ≤ 0.01. *** p ≤ 0.001. n = 4 (NU7441, ibrutinib, GSK8612), n = 5 (BX-795, sotrastaurin, PHA-665752), n = 8 (U73122, SB 743921 HCl, BIRB 796).

**Fig. S5:**
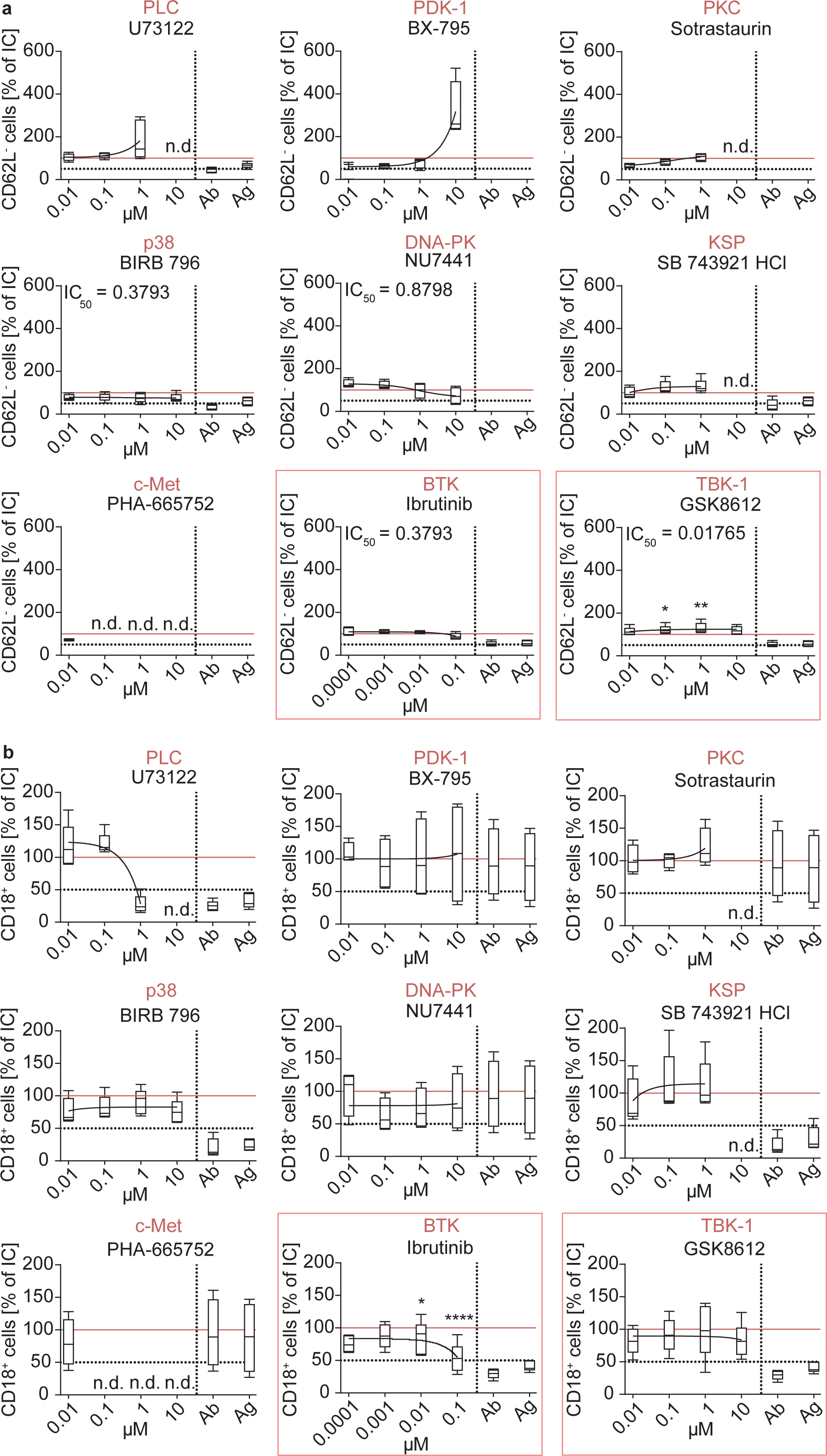
CD18 expression, but not CD62L shedding, is inhibited by ibrutinib. Neutrophils were stimulated with immobilized COL7-IC, treated with STIs for 2 h, and analyzed by flow cytometry for neutrophil activation markers. **(a)** CD18 expression **(b)** CD62L shedding. STIs that were chosen based on the multiplex kinase activity profiling are highlighted by red boxes. Kruskal-Wallis test with Dunn’s multiple comparisons test was performed. n = 5. * p ≤ 0.05, **** p ≤ 0.0001. n.d., not determined.

**Fig. S6:**
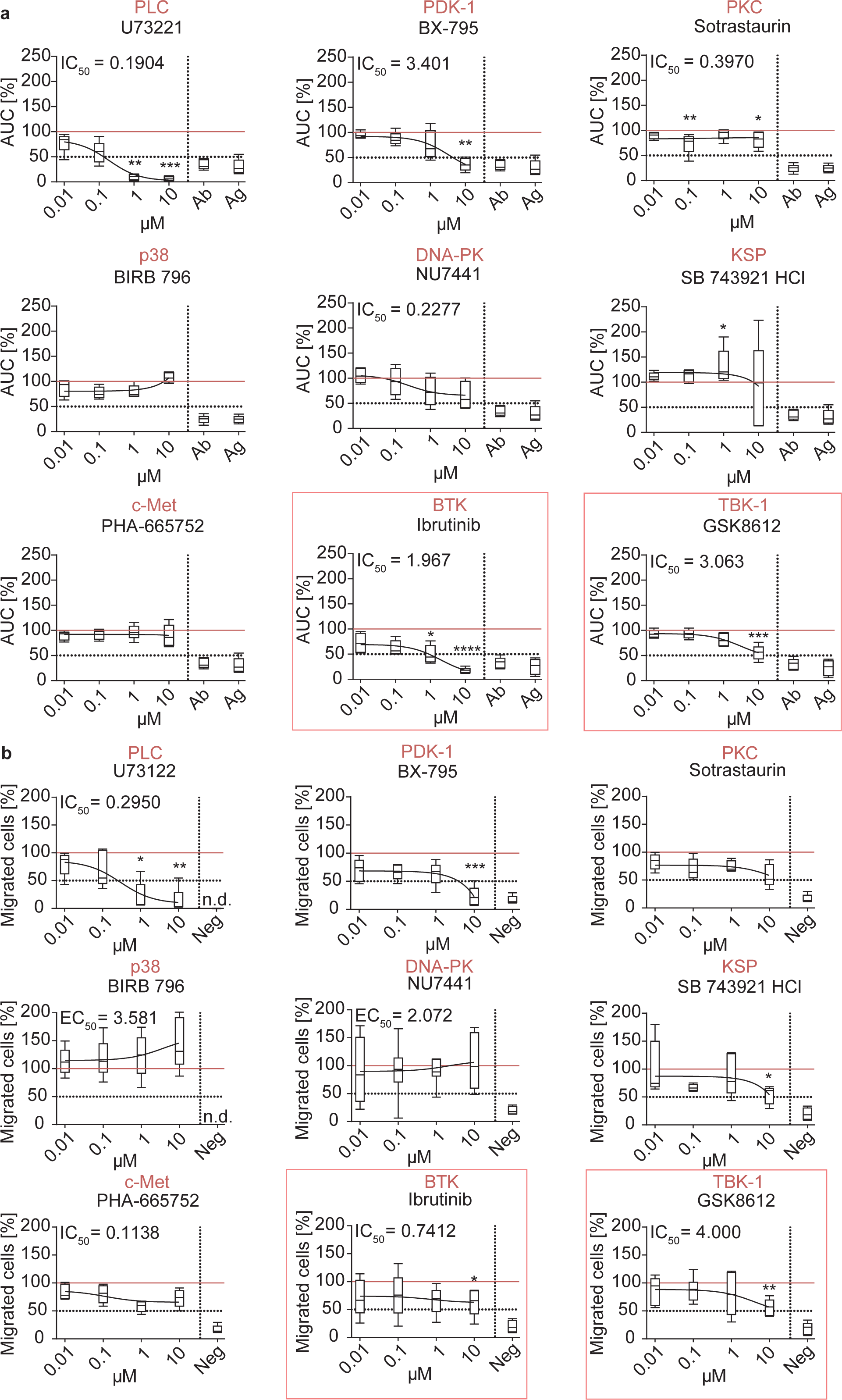
Several STIs impair neutrophil adhesion and chemotaxis. **(a)** Human neutrophils were activated with immobilized COL7-IC and treated with STIs for 2 h. **(b)** Human neutrophils treated with STIs were incubated in a Boyden chamber for 1 h to assess IL-8-mediated chemotaxis. STIs that were chosen based on the multiplex kinase activity profiling are highlighted by red boxes. Kruskal-Wallis test with Dunn’s multiple comparisons test was performed. n = 5. * p ≤ 0.05, ** p ≤ 0.01, *** p ≤ 0.001, **** p ≤ 0.0001.

**Fig. S7:**
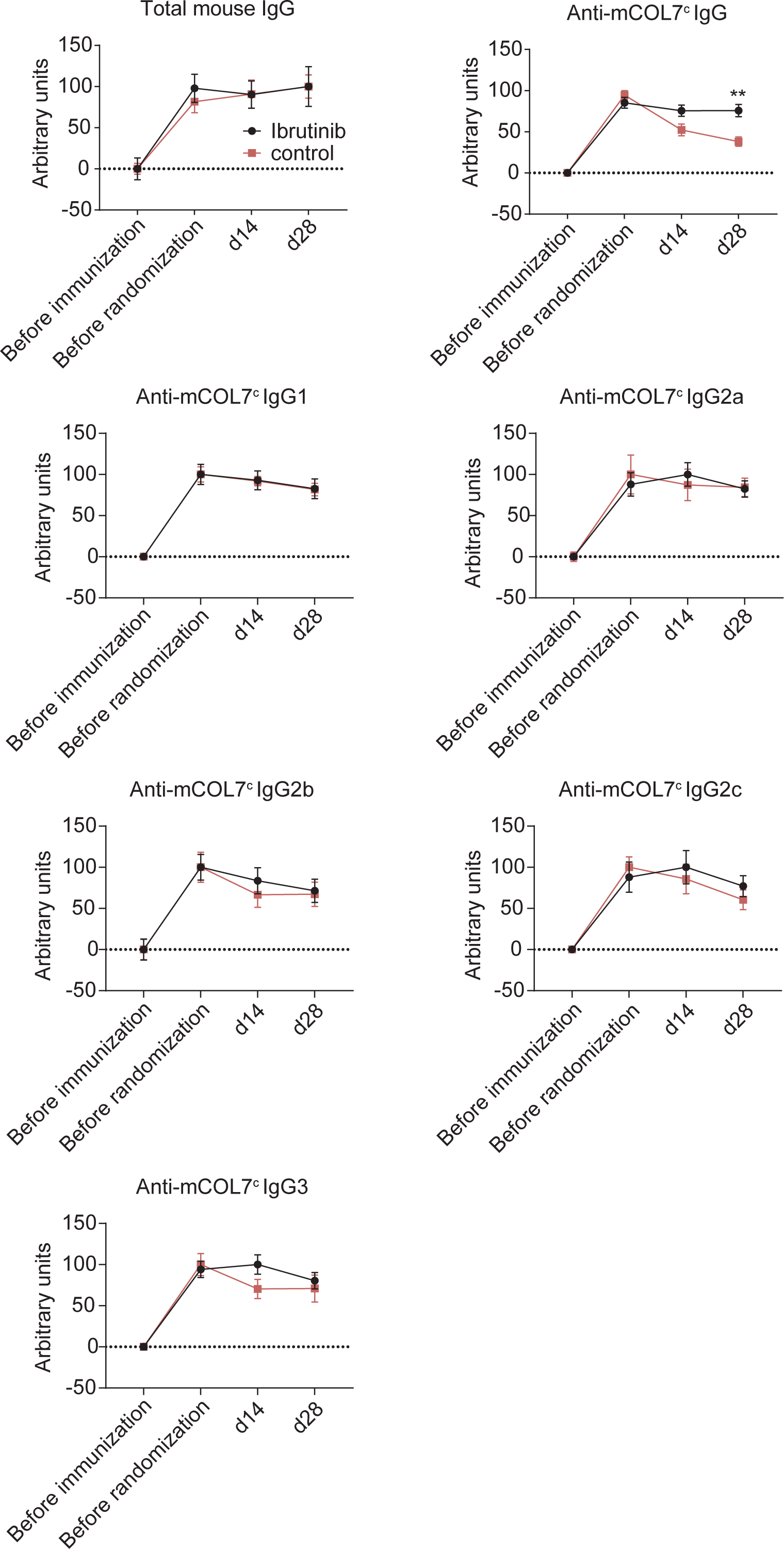
Ibrutinib does not alter total IgG titers or specific IgG subclass titers in active EBA. Antibody titers from serum of immunization-induced EBA mice and controls over the time course of the experiment. Total IgG, mCOL7^c^-specific IgG, and mCOL7^c^-IgG subclass concentrations were determined by ELISA. Values were normalized to the timepoint before immunization. Two-way ANOVA with Šidák’s multiple comparisons test was performed. n = 15. ** p ≤ 0.01.

**Fig. S8:**
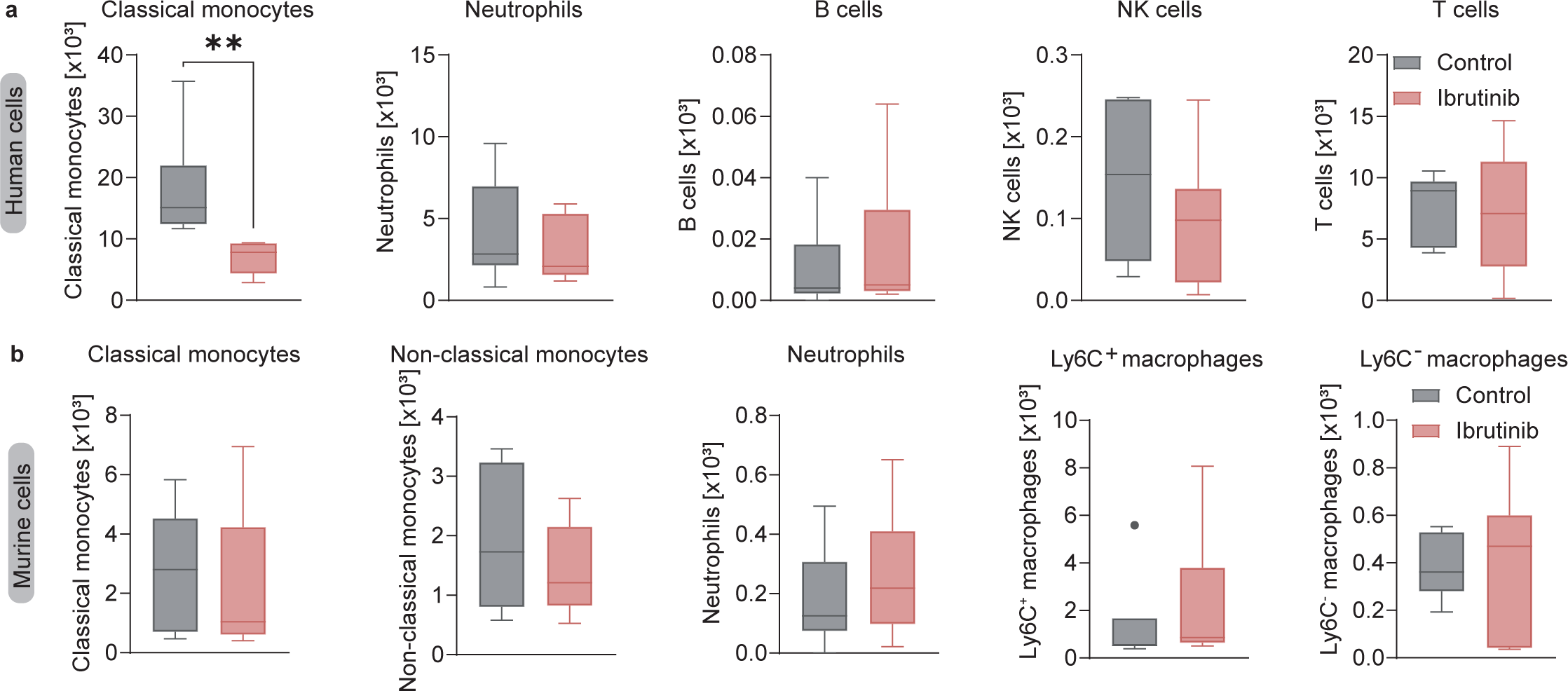
Less classical monocytes are detected in ibrutinib-treated HIS mice compared to control mice. EBA was induced in HIS mice by antibody transfer and the mice were treated with ibrutinib or vehicle for 12 days. Blood samples were taken on the final day to assess **(a)** human immune cell populations and **(b)** murine immune cell populations, by flow cytometry. Mann-Whitney test was conducted. n = 7 (control), n = 9 (ibrutinib). *** p ≤ 0.001.

**Fig. S9:**
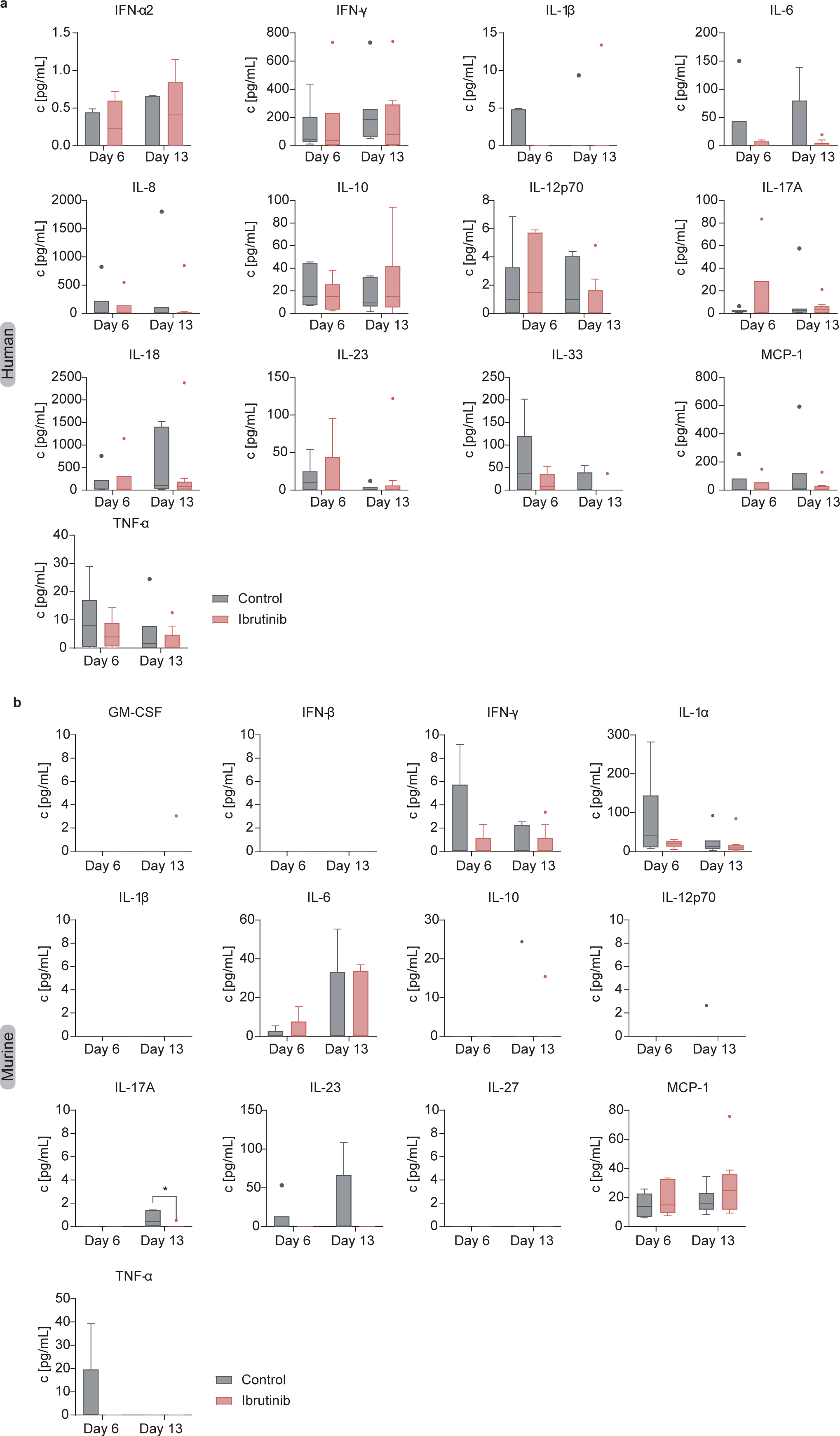
Murine cytokine expression is not changed to a biologically relevant extent. EBA was induced in HIS mice by antibody transfer and the mice were treated with ibrutinib or vehicle for 12 days. Blood samples were taken on day 6 and on the final day to assess **(a)** human and **(b)** murine serum cytokine levels. Mixed effects analysis with Šidák’s multiple comparisons test was performed. n = 7 (control), n = 9 (ibrutinib). *** p ≤ 0.001.

## Notes

https://www.ebi.ac.uk/ena/browser/view/PRJEB103987

### Competing Interest Statement

The authors have declared no competing interest.

### Summary of Updates

Minor changes have been made to the figures and the manuscript has been updated accordingly. In addition, the format of the manuscript has been adapted.

